# WASP restricts active Rac to maintain cells’ front-rear polarisation

**DOI:** 10.1101/602425

**Authors:** Clelia Amato, Peter Thomason, Andrew Davidson, Karthic Swaminathan, Shehab Ismail, Laura Machesky, Robert Insall

**Affiliations:** CRUK Beatson Institute, Switchback Road, Bearsden G61 1BD, UK; Institute of Cancer Sciences, University of Glasgow, University Avenue, Glasgow, G12 8QQ, UK

**Keywords:** Actin polymerization, Arp2/3 complex, Cell polarity, Uropod, Small GTPases, CRIB motif, Actin cytoskeleton

## Abstract

Efficient motility requires polarised cells, with anterior pseudopods and a retracting rear. This polarisation requires that the pseudopod catalyst Rac is restricted to the front. Here we show that the Arp2/3 complex regulator WASP is important for maintaining front–rear polarity, using a mechanism that limits where active Rac localises. *Dictyostelium* cells lacking WASP inappropriately activate Rac and SCAR/WAVE at their rears, leading to reduced cell speed. WASP facilitates the internalisation of clathrin-coated pits, and its Rac-binding CRIB motif is considered essential for its localisation and activity. However, WASP mutants with deleted CRIB domains, or harbouring a new mutation that prevents Rac binding, localise normally, recruit Arp2/3 complex, and drive actin polymerisation. Similarly, Rac inhibitors do not block WASP localisation or activation. Despite this, WASP CRIB mutants cannot restore polarisation of active Rac. Thus, WASP’s interaction with Rac regulates Rac activity and cell polarity, but is dispensable for activating actin polymerization.

## Introduction

Filamentous actin (F-actin) fulfils a number of key functions in migrating cells. One particularly important role is the generation of protrusions, known by a range of names such as pseudopods, lamellipods and filopods. A second is maintenance of efficient intracellular trafficking, by driving endocytosis and ensuring vesicle sorting. These functions depend upon the highly conserved Arp2/3 complex, which drives branching and growth of the F-actin network (Mullins et al., 1998).

Cells regulate the localisation and activity of the Arp2/3 complex using the WASP family of actin nucleation-promoting factors (NPFs). The founding member of the family, WASP, was discovered as the cause of the immune deficiency Wiskott-Aldrich syndrome (Derry et al., 1994). WASP has since been shown to be specific to haematopoietic lineages; WASP mutations that cause Wiskott-Aldrich syndrome consequently lead to limited and specific conditions such as microthrombocytopenia, rather than organism-wide effects on cell function. Vertebrates, however, possess a ubiquitous WASP paralogue. In mammals, this is termed N-WASP and was originally described as a neural-specific gene (Miki et al., 1996), although is now known to be expressed in nearly all cell types (Miki et al., 1998). Most eukaryotic cells, including *Dictyostelium*, yeasts and *Drosophila*, express a single WASP protein, which is the orthologue of vertebrates’ ubiquitous isoform (Veltman and Insall, 2010) although plants, and a subset of organisms from divergent clades, do not possess WASP genes. In most cases, the principal role of WASPs is to facilitate clathrin-mediated endocytosis (CME) (Davidson et al., 2018; Kochubey et al., 2006; Madania et al., 1999; Merrifield et al., 2004).

Other members of the WASP family recruit and activate the Arp2/3 complex to other structures and for other purposes. For instance, the SCAR/WAVE complex drives formation of actin protrusions such as pseudopods and lamellipods (Evans et al., 2013; Kunda et al., 2003; Veltman et al., 2012; Weiner et al., 2006), and is therefore a key regulator of cell migration and polarisation.

Protrusions can be generated in virtually every region of the plasma membrane, but those that cause locomotion are usually initiated and maintained at the front of the cell, where active Rac and SCAR/WAVE are localised. Whether clathrin-coated pits (CCPs) are internalised in a defined area of the cell appears to be cell type-dependent. The weight of data suggest that CME typically occurs at the rear half of highly motile cells, including lymphocytes (Samaniego et al., 2007), leukocytes (Davis et al., 1982), and *Dictyostelium* (Damer and O’Halloran, 2000).

How cells segregate WASPs and SCAR/WAVE, both spatially and functionally, is fascinating and not yet fully understood. Further complicating this issue, it has been shown in at least two evolutionarily distant organisms - *Dictyostelium* and *Caenorhabditis-* that WASP can compensate for loss of SCAR/WAVE (Veltman et al., 2012; Zhu et al., 2016), becoming promptly directed to sites of protrusions. This finding demonstrates that WASP is capable of responding to the upstream signals that drive pseudopod extension but is not normally employed to do so. To date, what determines the tight and yet plastic spatial and functional segregation of WASP and SCAR/WAVE remains to be elucidated.

One obvious prediction is that upstream regulators may have a crucial role. If so, localised accumulation of activators may account for the different subcellular distribution and function of the two NPFs. However, some of the upstream regulators (in particular active Rac) appear to be shared between WASPs and SCAR/WAVE. But while both genetics and cell biology consistently connect SCAR/WAVE to Rac1 (Chen et al., 2010; Eden et al., 2002; Rohn et al., 2011), there is no coherent model of how GTPases regulate WASPs. Most of our current knowledge on WASPs’ regulation arises from biochemical studies, which are often contradictory. WASPs from most organisms contain a CRIB domain, a short region that binds specifically to active small GTPases of the Rho family. CRIB domains are selective for Rac and Cdc42, rather than Rho itself (Burbelo et al., 1995). Early measurements of the ability of the ubiquitous N-WASP to trigger actin polymerisation *in vitro* concluded that Cdc42 had a major role (Miki et al., 1998; Rohatgi et al., 1999). However, more recent work suggests that Cdc42 alone specifically activates haematopoietic WASP, while Rac1 also interacts with N-WASP (Tomasevic et al., 2007). For *Dictyostelium* WASP, most attention has focussed on the unusual RacC (Han et al., 2006). However, the published data also shows WASP binding to the members of the Rac1 family (Han et al., 2006), which are expressed far more abundantly than RacC (data from Dictyexpress, https://dictyexpress.research.bcm.edu/bcm/), and are more closely related to Cdc42 and Racs from other organisms. The *Dictyostelium* genome does not contain a Cdc42 gene (Rivero et al., 2001), despite a number of Rac relatives. While Cdc42 and Rac1 appear to be crucial for WASPs to trigger actin polymerisation *in vitro*, their requirement for formation of actin filaments in living cells has been questioned. For instance, a CRIB-deleted WASP is able to rescue phenotypes of *Drosophila* mutants (Tal et al., 2002). Of interest, yeasts’ WASP (which also mediates CCPs internalisation), is devoid of a CRIB motif, raising the question as to how the NPF becomes activated during CME. To date there is no clear explanation regarding the inconsistency between the *in vitro* and *in vivo* data, nor is it fully understood whether WASP relies on a direct interaction with active GTPases to fulfil any of its cellular functions. In particular, the importance of Rac and Cdc42 for WASP activation during clathrin-mediated endocytosis remains substantially unexplored.

The requirement of small GTPases for activation of WASPs is a fascinating issue *per se*. However, it becomes an even more urgent point to clarify in order to understand how cells achieve spatial and functional segregation of WASPs and SCAR/WAVE. Indeed, a model whereby Rac mediates the activation of both NPFs fits poorly with the hypothesis that upstream regulators are responsible for their distinct sub-cellular localisation and functionality.

Recent work from our lab offers a fresh perspective on how cells may maintain spatial and functional separation of WASP and SCAR/WAVE. Loss of WASP (unlike loss of SCAR/WAVE) causes an unusual rear retraction defect in *Dictyostelium* cells. Further analysis shows that this is caused by aberrant accumulation of SCAR/WAVE at the back of migrating cells. This role for WASP in maintenance of front–rear polarity is intriguing and unforeseen, and does not fit well with existing models of cell polarity. In this study, we dissect the role of WASP in cell polarity, and clarify the importance of small GTPases for WASP function.

In summary, we uncover an unexpected role for WASP in maintenance of front–rear polarity, which works by restricting the regions of the cell that accumulate active Rac1. Furthermore, this work demonstrates the dispensability of direct interaction with active GTPases for WASP to form F-actin during clathrin-mediated endocytosis. Rac binding is still clearly important, for example in allowing WASP to make pseudopods in SCAR/WAVE’s absence. More provocatively, our study suggests an additional, reversed role for the interaction between WASP and GTPases – the presence of a CRIB motif may not only mean that WASP activity requires GTPase regulation, but that WASP modulates the distribution of GTPases after they are activated.

## Results

### Loss of WASP causes accumulation of SCAR/WAVE and active Rac at the rear

Previous work shows that knockout mutants in the *Dictyostelium* gene encoding WASP, *wasA*, migrate slower than their wild type counterparts, and that this is not due to a reduced rate of pseudopod extension, but rather to a defect in rear retraction (Davidson et al., 2018). More detailed analysis revealed that while wild type cells confine SCAR/WAVE at the extending protrusions (Fig. 1A), *wasA*^-^ cells also accumulate SCAR/WAVE within an enlarged rear uropod (Fig. 1B). Although able to maintain a morphologically dominant front, *wasA*^-^ cells fail to confine the leading edge-specific NPF to a single region of their plasma membrane, resulting in a nearly-bipolar appearance. We examined the drivers of *wasA*^-^ cells’ inability to exclude SCAR/WAVE from their rear. One intriguing possibility was that WASP may be responsible for the spatial confinement of SCAR/WAVE’s activators, therefore preventing the initiation of an actin polymerisation cascade in unwanted areas of the cell. Since active Rac is known as one of the major regulators of SCAR/WAVE (Chen et al., 2010; Eden et al., 2002; Lebensohn and Kirschner, 2009), we asked whether the localisation or dynamics of the GTP-bound GTPase was affected in *wasA*^-^ cells.

**Figure 1.**
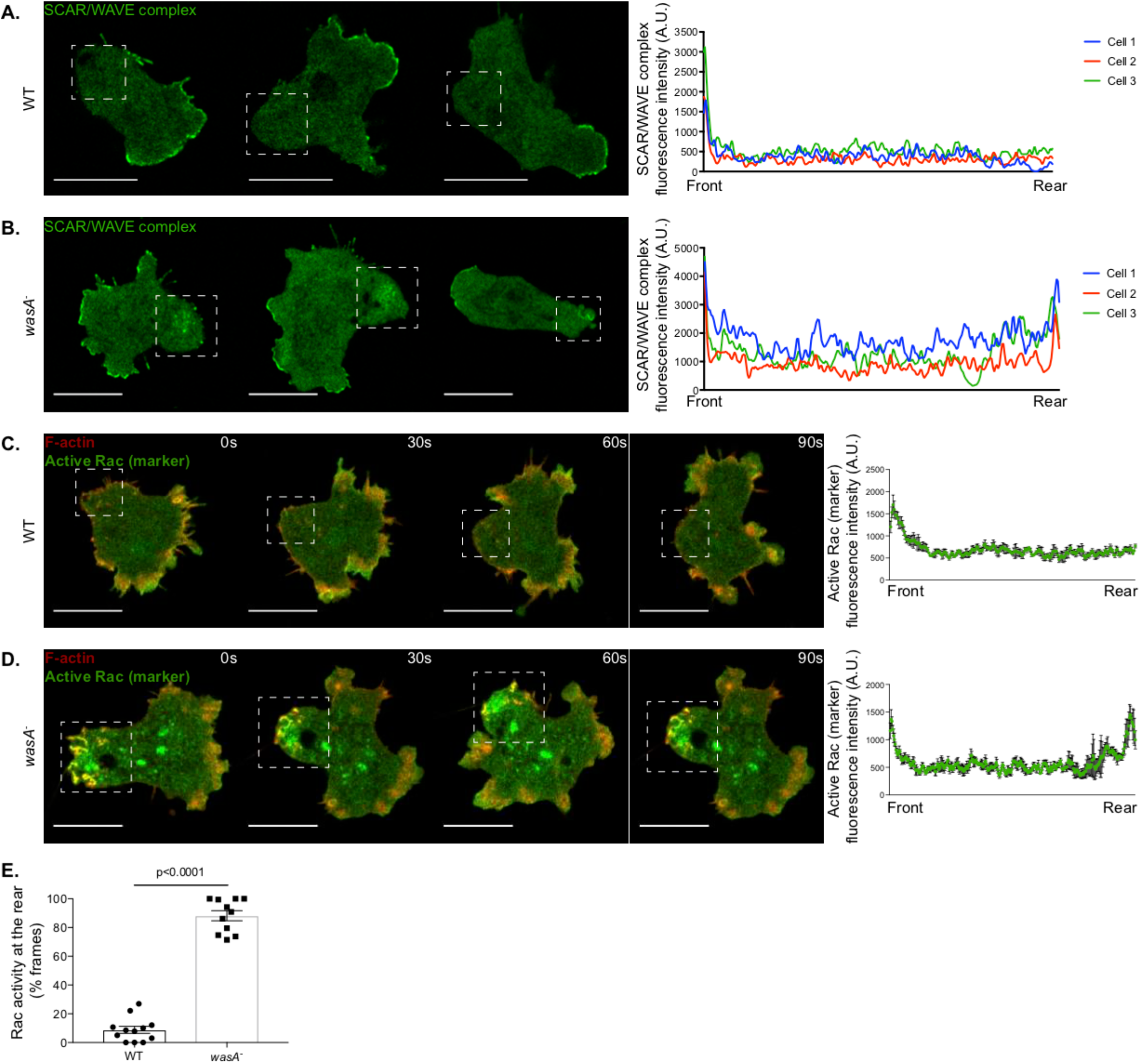
Migrating WASP null cells accumulate SCAR/WAVE and active Rac at the rear. **A and B.** The SCAR/WAVE complex (HSPC300-GFP) is confined to the front and excluded from the rear (dashed squares) of migrating wild type cells (WT) **(A)**, while is present both at the front and rear (dashed squares) of WASP null cells (*wasA*^-^) **(B)**. Fluorescence plot reports the intensity of the SCAR/WAVE complex along a straight line connecting the front to the rear of three representative cells at a given time point. Scale bars represent 10 μm. **C and D.** Active Rac (PakB CRIB-GFP) is confined to the front and excluded from the rear (dashed squares) of a migrating WT cell **(C)**, but is present at the leading edge as well as at the enlarged uropod (dashed squares) of a *wasA*^-^ cell **(D)**. Fluorescence plots quantify the active Rac enrichment along a straight line connecting the front to the rear of individual cells at four time points. F-actin (LifeAct-mRFP) can be detected at the front as well as on transient spots at the rear half (dashed squares) of a WT cell **(C)**, and at the front but also on persistent structures at the rear of a *wasA*^-^ cell **(D)**. Scale bars represent 10 μm. **E.** Frequency of active Rac enrichment at the rear half of migrating wild type (WT) and *wasA*^-^ cells. Wild type cells (n=12 cells): 8.9 ± 2.5 % frames; *wasA*^-^ cells (n=11 cells): 88.2 ± 3.5 % frames; means ± SEM. Mann Whitney test, p <0.0001.

We performed live-cell imaging on *wasA* knockout and wild type cells expressing the isolated CRIB motif of PakB, which binds to Racs with high affinity *in vitro* (de la Roche et al., 2005), and has been previously described as a reporter for active Rac in *Dictyostelium* (Veltman et al., 2012). Similar constructs, mostly based on the Rac-binding motif of PAK1, have been used to monitor the dynamics of endogenous active Rac in living mammalian cells (Kraynov et al., 2000; Weiner et al., 2007; Xu et al., 2003). As expected, migrating wild type cells accumulate active Rac at the leading edge (Fig. 1C and S1A), where it co-localises with F-actin (as shown by GFP-LifeAct). However, *wasA* knockout cells accumulate active Rac both at the front and at the enlarged rear (Fig. 1D and S1B), leading to the formation of F-actin-rich structures at both ends during migration. Detailed quantification revealed that while wild type cells accumulate active Rac at the rear in a small number of frames (most of which coincide with pseudopods that are swept from the front to the back of the cell as it moves), *wasA* knockout cells accumulate active Rac at their back in a significantly higher percentage of frames (Fig. 1E). Since the localisation of both active Rac and SCAR/WAVE is aberrant in *wasA* knockout cells, and given that Rac directly regulates SCAR/WAVE, the simplest conclusion would be that WASP limits the amount of membrane with active Rac.

### Generation of WASP CRIB mutants

We tested possible molecular mechanisms underlying WASP-mediated confinement of active Rac and SCAR/WAVE. We started by focusing on WASP’s CRIB (Cdc42/Rac interacting binding) motif, a relatively short stretch that has been found in a number of functionally-unrelated proteins and in evolutionarily-distant contexts (Burbelo et al., 1995; Pirone et al., 2001). The CRIB motif has long been considered a marker of direct binding to small GTPases, typically seen in effectors of Rac1 or Cdc42. Accordingly, WASP has been proposed to act downstream of Rac1 and/or Cdc42 mostly based on *in vitro* data (Miki et al., 1998; Rohatgi et al., 1999; Tomasevic et al., 2007). However, it remains mysterious what the physiological connection between small GTPase activity and WASP activation may be.

To analyse the importance of the interaction between WASP and active Rac, we generated WASPs that could not interact with Rac by introducing mutations in the CRIB motif (Fig. 2A). In the first, WASP^ΔCRIB^, we deleted 14 residues (173-186) that constitute a large proportion of the CRIB’s core (Burbelo et al., 1995). A related mutant has been described in *Drosophila* (Tal et al., 2002). We were concerned that this substantial deletion could affect protein function, so we designed a second mutant, hereafter referred to as WASP^**CRIB^, containing only two conservative amino acidic changes (I173A; F179A). These residues were chosen based on their inferred position in the WASP/Rac interface (Fig. 2B), based on the structure of the complex between Cdc42 and the minimal PBD (p21 binding domain, which includes the CRIB motif) of WASP (Abdul-Manan et al., 1999). The two residues are well conserved, and occupy dedicated pockets within the interface between GTPase and CRIB; changing them to different hydrophobic amino acids is expected to maximally diminish the binding energy, with minimal change to the CRIB motif’s structure. Importantly, both changes only affect the N-terminus of the CRIB motif, which does not appear to be primarily involved in maintenance of the autoinhibited conformation (Kim et al., 2000). We therefore do not expect mutating the N-terminus of the CRIB motif to steer WASP to an open conformation or render it inappropriately active.

**Figure 2.**
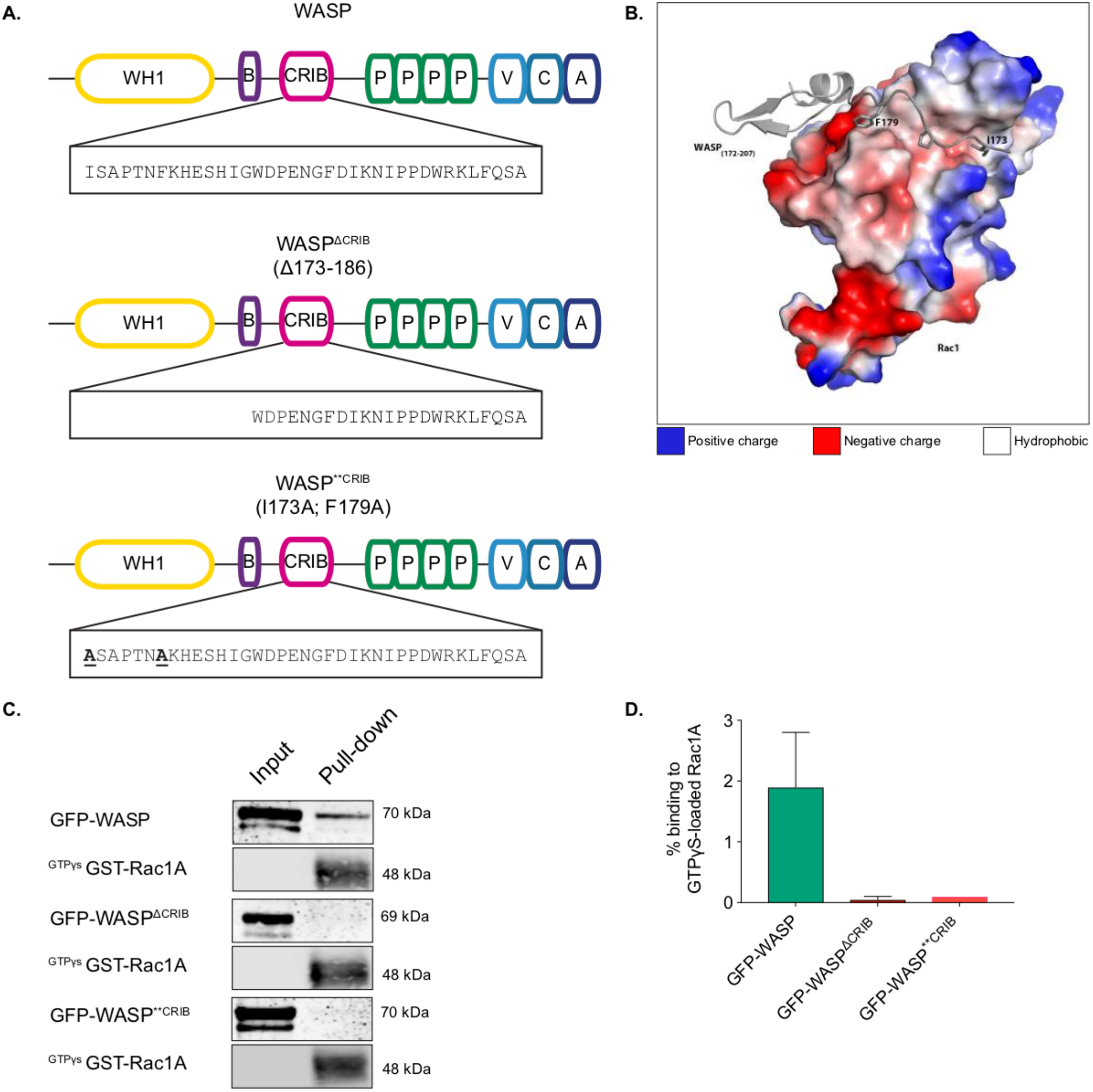
Mutations in the WASP CRIB motif abrogate binding to active Rac1. **A.** Schematic representation of WASP domain composition and mutations introduced within the CRIB motif. Top panel: wild type WASP. Middle panel: WASP^ΔCRIB^, harbouring a deletion of 14 residues (Δ173-186) at the N-terminus of the CRIB motif. Bottom panel: WASP^**CRIB^, harbouring two point mutations at the N-terminus of the CRIB motif. **B.** 3D representation of the WASP / Rac1 interface. WASP residues I173 and F179 (in grey) establish contacts with a hydrophobic (white) region of Rac1. **C.** Immunoblot showing that GFP-WASP (first panel) interacts with active (GTPγs-bound) Rac1A, while GFP-WASP^ΔCRIB^ and GFP-WASP^**CRIB^ (third and fifth panels respectively) do not (IB= anti-GFP). Anti-GST western blot was performed (second, fourth and sixth panels) to verify the expression of GST-Rac1A. **D.** Immunoblot quantification showing that no binding of GFP-WASP^ΔCRIB^ and GFP-WASP^**CRIB^ to active Rac1 occurs (GFP-WASP: 1.9% ± 0,9; GFP-WASP^ΔCRIB^: 0.05% ± 0.05, GFP-WASP^**CRIB^: 0.1, mean ± SEM, n= 3).

We tested the ability of the two WASP CRIB mutants to interact with active Racs using pull-down assays. We examined Rac1A, the most highly expressed (data from Dictyexpress, https://dictyexpress.research.bcm.edu/bcm/) and most closely related to canonical Rac1 of other species, and RacC, which has been specifically associated with *Dictyostelium* WASP (Han et al., 2006). Bacterially-purified GST-Racs were activated by loading with GTPγs, a non-hydrolysable analogue of GTP. As shown in figure 2C and D and supplementary figure 2, wild type WASP binds to active Racs, but neither CRIB mutant is able to bind to either GTPase.

### Rac binding is not required to recruit WASP to clathrin coated pits

We first assessed the functionality of the WASP CRIB mutants.

The most widely-accepted model of WASP regulation asserts that Rac or Cdc42 have an essential role in controlling WASP’s activity (Han et al., 2006; Miki et al., 1998; Rohatgi et al., 1999; Tomasevic et al., 2007). According to this model, the inability of the WASP CRIB mutants to interact with active Rac should obstruct WASP function. We therefore tested it by live-cell imaging of *wasA*^-^ cells co-expressing fluorescently tagged WASP mutants and clathrin. This approach allowed us to assess whether a functional CRIB domain was important for WASP’s biochemical activity, and more generally to test which physiological functions of WASP depend on Rac regulation.

As shown in figure 3A and B, wild type WASP generates puncta that overlap with CCPs. At any single time, the number of clathrin puncta that are also WASP-positive is low due to the fact that WASP only appears as a brief burst, while CCPs persist on the plasma membrane for a longer period. Figure 3C shows a representative example of clathrin/WASP dynamics visualised using TIRF (Total Internal Reflection Fluorescence) microscopy of cells gently compressed under agarose, which diminishes front-rear polarity. WASP is recruited (t=0 s) on a pre-existing CCP, where it sits for a few seconds before disappearing along with clathrin. Clearance of WASP and clathrin from the TIRF field is considered as a *bona fide* sign of CCP internalisation (Merrifield et al., 2002).

**Figure 3.**
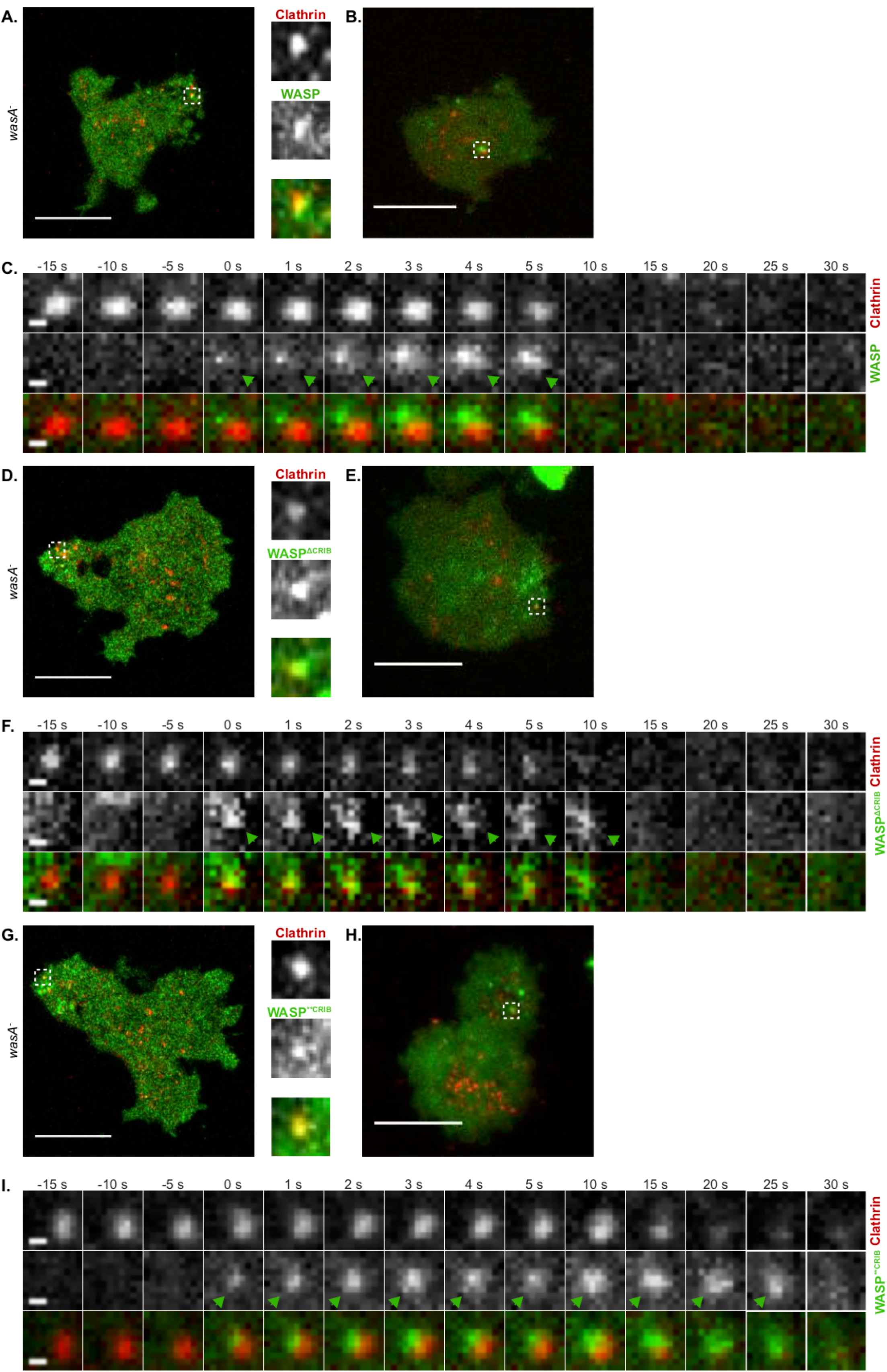
WASP does not require a direct interaction with active Rac to be recruited to clathrin-coated pits. **A.** Airyscan confocal microscopy of a migrating WASP null cell (*wasA*^-^) co-expressing WASP (GFP-WASP, rescue) and clathrin (clathrin light chain-mRFPmars). The punctum highlighted by the dashed square, shown in greater detail within the insets, is an example of clathrin/WASP co-localisation. Scale bar represents 10 μm. **B.** TIRF microscopy of a *wasA*^-^ cell co-expressing GFP-WASP and clathrin light chain-mRFPmars gently compressed under agarose. An example of WASP and clathrin co-localisation is highlighted by the dashed square. Scale bar represents 10 μm. **C.** WASP recruitment on a clathrin-coated pit. GFP-WASP appears (t= 0 s, green arrowhead) on a pre-existing clathrin spot; after a short time, clathrin and WASP disappear synchronously from the TIRF field (t= 10 s). Scale bars represent 0.5 μm. **D.** Airyscan confocal microscopy of a migrating *wasA*^-^ cell co-expressing GFP-WASP^ΔCRIB^ and clathrin light chain-mRFPmars. An example of co-localisation is highlighted by the dashed square and shown in greater detail within the insets. Scale bars represent 10 μm. **E.** TIRF microscopy of a *wasA*^-^ cell co-expressing GFP-WASP^ΔCRIB^ and clathrin light chain-mRFPmars gently compressed under agarose. The punctum highlighted by the dashed square shows an example of co-localisation between clathrin and WASP^ΔCRIB^. Scale bar represents 10 μm. **F.** WASP^ΔCRIB^ recruitment on a clathrin-coated pit. GFP-WASP^ΔCRIB^ appears on a pre-existing clathrin punctum (t= 0 s, green arrowhead), where it persists for a longer period of time in comparison to wild type WASP. Ultimately, clathrin and GFP-WASP^ΔCRIB^ leave the TIRF field (t= 15 s). Scale bars represent 0.5 μm. **G.** Airyscan confocal microscopy of a migrating *wasA*^-^ cell co-expressing GFP-WASP^**CRIB^ and clathrin light chain-mRFPmars. An example of co-localisation is highlighted by the dashed square and shown in greater detail within the insets. Scale bars represent 10 μm. **H.** TIRF microscopy of a *wasA*^-^ cell co-expressing GFP-WASP^**CRIB^ and clathrin light chain-mRFPmars gently compressed under agarose. The punctum highlighted by the dashed square shows an example of co-localisation. Scale bar represents 10 μm. **I.** WASP^**CRIB^ recruitment on a clathrin-coated pit. GFP-WASP^**CRIB^ appears on a pre-existing clathrin punctum (t= 0 s, green arrowhead), where it remains for an extended period of time in comparison to wild type WASP. Eventually, both clathrin and GFP-WASP^**CRIB^ disappear from the TIRF field (t= 25 s). Scale bars represent 0.5 μm.

To our surprise, we found that both WASP CRIB mutants generate puncta that coincide with CCPs (figures 3D, E, and G, H). Live TIRF imaging confirmed that WASP^ΔCRIB^ and WASP^**CRIB^ are recruited to pre-existing clathrin puncta (t= 0 s in figures 3F and I).

This clearly demonstrates that a direct interaction with active Rac is not required for WASP to be recruited to clathrin-coated pits, suggesting that factors other than GTPases control WASP’s sub-cellular localisation in living cells during endocytosis.

### WASP does not require Rac to recruit the Arp2/3 complex and actin

We noticed that while spots generated by wild type WASP tend to last only a few seconds, those generated by the two WASP CRIB mutants co-localise with CCPs for a longer time. One possible cause underlying a delay/impairment in CME could be the lack of actin polymerisation on CCPs, which is mandatory in yeast and *Dictyostelium* (Davidson et al., 2018; Kaksonen et al., 2003), but not in all mammalian cells and not in all membranes from the same cell type (Fujimoto et al., 2000; Gottlieb et al., 1993). Mechanistically, nucleation of actin filaments at sites of CCPs internalisation is thought to provide the force required to facilitate membrane invagination (Aghamohammadzadeh and Ayscough, 2009; Boulant et al., 2011). Given our previous report showing that loss of WASP causes an increase in CCP lifetime due to the lack of actin polymerisation [Fig. 2E; (Davidson et al., 2018)], we tested whether WASP CRIB mutants are able to trigger actin polymerisation once recruited to CCPs, using live imaging of *wasA*^-^ cells co-expressing GFP-WASP and RFP-tagged Arp2/3 complex. As shown in Figure 4A, WASP puncta tend to be evenly distributed at the rear half of polarised cells. At essentially all times observed, WASP spots are Arp2/3 complex-positive; this presumably reflects the fact that WASP is activated as it is targeted to clathrin spots, so the Arp2/3 complex is recruited synchronously. This is clearly shown in figure 4B, which reports a typical example of WASP and Arp2/3 complex recruitment.

**Figure 4.**
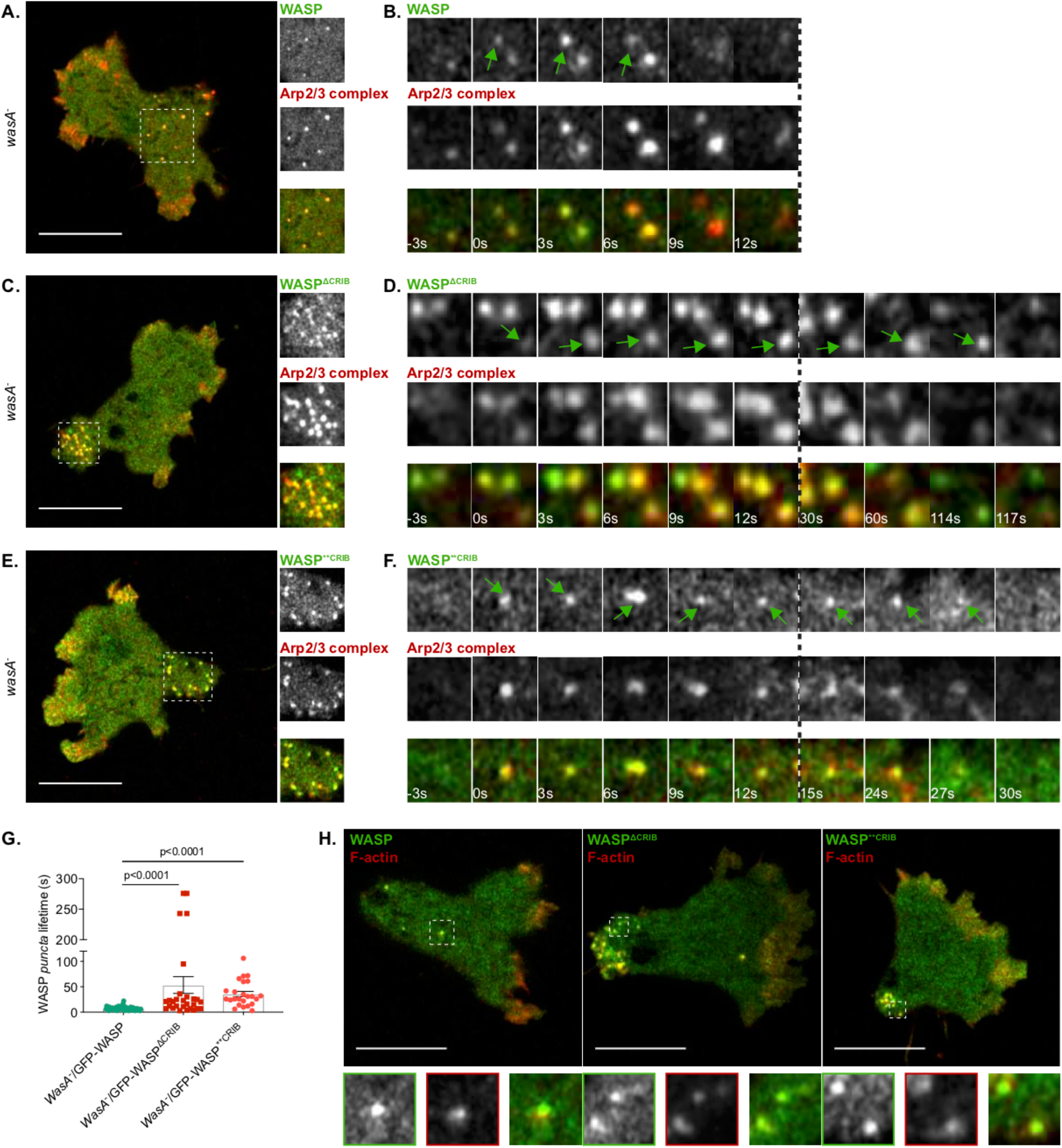
WASP does not require a direct interaction with active Rac to recruit the Arp2/3 complex and trigger actin polymerisation on puncta. **A.** Airyscan confocal imaging of a migrating WASP null cell (*wasA*^-^) co-expressing GFP-WASP (rescue) and mRFPmars2-ArpC4. WASP puncta are evenly distributed at the rear of the cell and are all enriched in Arp2/3 complex. The area highlighted by the dashed square is shown in greater detail within the insets. Scale bar represents 10 μm. **B.** WASP and Arp2/3 complex recruitment on puncta. GFP-WASP appears on a punctum (t= 0 s, green arrow) alongside mRFPmars2-ArpC4; both last for a short period time and synchronously disappear (t= 9 s in the example shown). **C.** Airyscan confocal imaging of a migrating *wasA*^-^ co-expressing GFP-WASP^ΔCRIB^ and mRFPmars2-ArpC4. All WASP^ΔCRIB^ puncta cluster within the enlarged uropod (dashed square). As shown in greater detail within the inset, all GFP-WASP^ΔCRIB^ spots are also enriched in Arp2/3 complex. Scale bar represents 10 μm. **D.** WASP^ΔCRIB^ and Arp2/3 complex recruitment on puncta. GFP-WASP^ΔCRIB^ appears as a punctum (t= 0 s, green arrow) along with mRFPmars2-ArpC4. WASP^ΔCRIB^ remains within spots (alongside the Arp2/3 complex) for a longer period of time than its wild type counterpart (t= 117 s in the example shown). **E.** Airyscan confocal imaging of migrating *wasA*^-^ cells co-expressing GFP-WASP^**CRIB^ and mRFPmars2-ArpC4. All WASP^**CRIB^ puncta cluster within the enlarged rear (dashed square). As shown in the inset, all WASP^ΔCRIB^ spots are also Arp2/3 complex-positive. Scale bar represents 10 μm. **F.** WASP^**CRIB^ and Arp2/3 complex recruitment on puncta. GFP-WASP^**CRIB^ appears as a spot (t= 0 s, green arrow) alongside mRFPmars2-ArpC4. WASP^**CRIB^ (as well as the Arp2/3 complex) remains within the punctum for a longer period of time than wild type WASP (t= 27 s in the example shown). **G.** Lifetime of wild type and WASP CRIB-mutants puncta. *wasA*^-^/GFP-WASP (n= 52 puncta): 7.2 s ± 0.5; *wasA*^-^/GFP-WASP^ΔCRIB^ (n= 28 puncta): 53.4 s ± 16.6; *wasA*^-^/GFP-WASP^**CRIB^ (n= 24 puncta): 35.8 s ± 5.1; means ± SEM. Kruskal-Wallis test, *wasA*^-^/GFP-WASP vs *wasA*^-^ /GFP-WASP^ΔCRIB^ = p<0.0001; *wasA*^-^/GFP-WASP vs *wasA*^-^/GFP-WASP^**CRIB^= p <0.0001). **H.** Airyscan confocal imaging of migrating *wasA*^-^ cells expressing LifeAct-mRFPmars2 along with GFP-WASP (rescue, left panel), GFP-WASP^ΔCRIB^ (central panel), GFP-WASP^**CRIB^ (right panel). All puncta generated by wild type WASP or WASP CRIB mutants are enriched in actin filaments. Examples highlighted by dashed squares are shown in greater detail in the insets. Scale bars represent 10 μm.

Unexpectedly in the light of our current understanding of WASP activation, WASP^ΔCRIB^ and WASP^**CRIB^ are both fully able to recruit the Arp2/3 complex to puncta. As seen with wild type WASP, all puncta generated by either of the CRIB-mutated WASPs are also Arp2/3 complex-positive (Figures 4C, D and E, F). Spots generated by WASP^ΔCRIB^ and WASP^**CRIB^ were more likely to be seen within the enlarged uropod, and appeared to show increased lifetimes, as reported in figure 4G. Each of the mutants shows a bimodal distribution, with many puncta internalised at a roughly normal rate, but a number lasting for several minutes.

To further confirm that actin is being polymerised on puncta generated by WASP CRIB mutants, we imaged *wasA*^-^ cells co-expressing GFP-WASP mutants and RFP-LifeAct (Figure 4H). Consistent with the Arp2/3 complex data, F-actin was observed on all WASP puncta irrespective of the presence of a functional CRIB domain, and therefore independently of small GTPase binding.

### Rac inhibition does not affect the ability of WASP to generate dynamic puncta

The ability of WASP to localise to CCPs and trigger Arp2/3 complex-mediated actin polymerisation without interacting with GTPases disagrees with most accepted models of WASP activation (Miki et al., 1998; Rohatgi et al., 1999; Tomasevic et al., 2007). For further experimental support, we therefore examined the dynamics of normal WASP in wild type cells upon chemical inhibition of Rac.

Several commercially available Rac inhibitors are functional and widely used in mammalian cells. However, no Rac inhibitor has yet been verified for *Dictyostelium*. To identify an effective Rac inhibitor, we transfected wild type cells with an active Rac marker and monitored its dynamics upon treatment with candidates. The first molecule tested, NSC23766 (Gao et al., 2004) failed to show any effect on *Dictyostelium* even when added at significantly higher concentrations than usual (data not shown). However, a second compound, EHT1864, which binds directly to Rac and blocks its loading with GTP and thus activation (Onesto et al., 2008; Shutes et al., 2007) was effective at a minimal dose of 3 μM. As shown in figure 5A, the active Rac marker is localised in discrete regions of the plasma membrane (mostly macropinosomes) prior to addition of the Rac inhibitor (t= −10 s). Low doses (0.3 μM) of the inhibitor cause no obvious changes in cell shape nor alteration of the active Rac marker localisation. However, higher concentrations (3 μM – 30 μM) cause rapid and complete loss of membrane localisation of the active Rac marker; cells consequently lose all ability to generate either macropinosomes or pseudopods, and rapidly become rounded. Thus, EHT1864 is an effective inhibitor of *Dictyostelium* Rac signalling.

**Figure 5.**
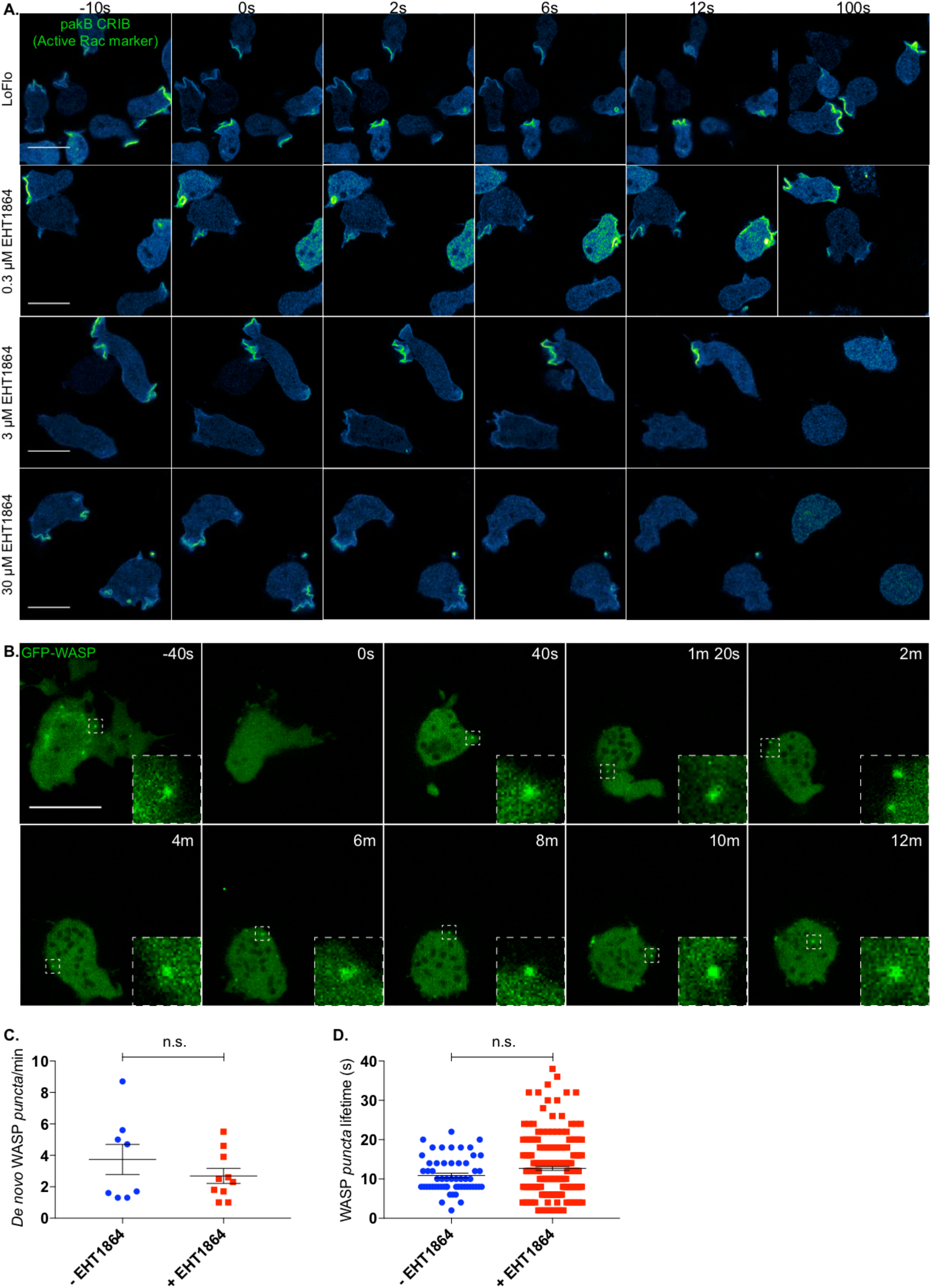
Rac inhibition does not measurably alter WASP dynamics. **A.** EHT1864 efficiently inhibits active Rac in *Dictyostelium*. Wild type cells expressing an active Rac marker (PakB CRIB-mRFPmars2) were imaged prior to and upon addition of increasing concentration of the Rac inhibitor EHT1864. Before treatment (t= −10 s), the active Rac marker is enriched on the plasma membrane, mostly at forming macropinosomes. Addition (t= 0 s) of low doses (0.3 μM) of the inhibitor does not affect cell morphology nor the distribution of the active Rac marker. Addition (t= 0 s) of a higher dose (3 μM) of the compound triggers a change in cell shape as well as a reduction of membrane-targeted active Rac marker. Further increase of the drug concentration (30 μM) leads to a quicker and more dramatic effect. Scale bars represent 10 μm. **B.** EHT1864 does not inhibit WASP punctate localisation. WASP dynamics upon chemical inhibition of Rac was investigated using Airyscan confocal imaging of WASP null cells (*wasA*^-^) expressing GFP-WASP. Before addition of the drug (t= −40 s), cells are polarised and WASP can be detected on individual spots (dashed square). Shortly after treatment (t= 0 s) with a high dose of the drug (50 μM), cells undergo a morphological change and ultimately round up. As highlighted by dashed squares at given time points, WASP is able to generate puncta even after 12 minutes from the treatment. Scale bar represents 10 μm. **C.** No significant difference can be measured between the number of WASP puncta generated per minute prior to and upon Rac inhibition. Untreated cells (-EHT1864, n= 8 cells): 3.7 ± 1.0; treated cells (+ EHT1864, n= 10 cells): 2.7 ± 0.5; means ± SEM. Unpaired t-test, p = 0.3130). **D.** No significant difference can be measured between the lifetime of WASP puncta generated prior to and upon addition of the Rac inhibitor. Untreated cells (-EHT1864, n= 57 puncta): 10.88 s ± 0.59; treated cells (+EHT1864, n= 213 puncta): 12.67 s ± 0.49; means ± SEM. Mann Whitney test, p= 0.2023.

We monitored the dynamics of GFP-WASP upon treatment with a high dose of EHT1864. As shown in figure 5B, cells undergo an immediate and dramatic morphological change once the compound has been added. However, in agreement with our data highlighting the ability of CRIB-mutated WASP to be recruited to CCP, we found that WASP is able to generate puncta up to 12 minutes after Rac inhibition. After this time, cells are generally so severely compromised that they tend to detach from the substrate and disappear from the field of view. Quantitative analysis (Figs 5C, D) shows that Rac inhibition by EHT1864 does not change the rate at which WASP puncta are generated. Similarly, the lifetimes of WASP spots generated before and after EHT1864 addition were not obviously different.

This result reinforces our conclusion that WASP does not need active GTPases to localise correctly during clathrin-mediated endocytosis, or to recruit Arp2/3 complex and generate F-actin once localised.

### WASP interaction with Rac is essential for front–rear polarity

The generation of WASP CRIB mutants allowed us to test the hypothesis that WASP may rely on a direct interaction with active Rac to maintain front–rear polarity during migration. To address this question, we imaged cells co-expressing mutant WASPs and active Rac marker during directed migration towards a source of chemoattractant.

As shown in figure 6A, and as reported in figure 1D, *wasA*^-^ cells recruit active Rac to the leading edge but also within the enlarged uropod. *wasA*^-^ cells expressing GFP-tagged wild type WASP behave normally, accumulating active Rac at the front, where pseudopods are being generated. The aberrant localisation of active Rac at the backs of cells is rescued (Fig. 6B). However, CRIB-mutated WASPs cannot suppress the accumulation of active Rac at the back of the cells (Fig. 6C and D). Cells expressing WASP CRIB mutants show an enlarged uropod, like that seen in unrescued *wasA*^-^ cells. Representative plots showing active Rac marker localisation from front to rear are shown in Fig.6E. Detailed quantification (reported in Fig. 6F) reveals that cells expressing CRIB-mutated WASPs accumulate active Rac at the rear to the same degree as *wasA*^-^ cells, while wild type WASP rescues the spatial confinement of active Rac at the leading edge.

**Figure 6.**
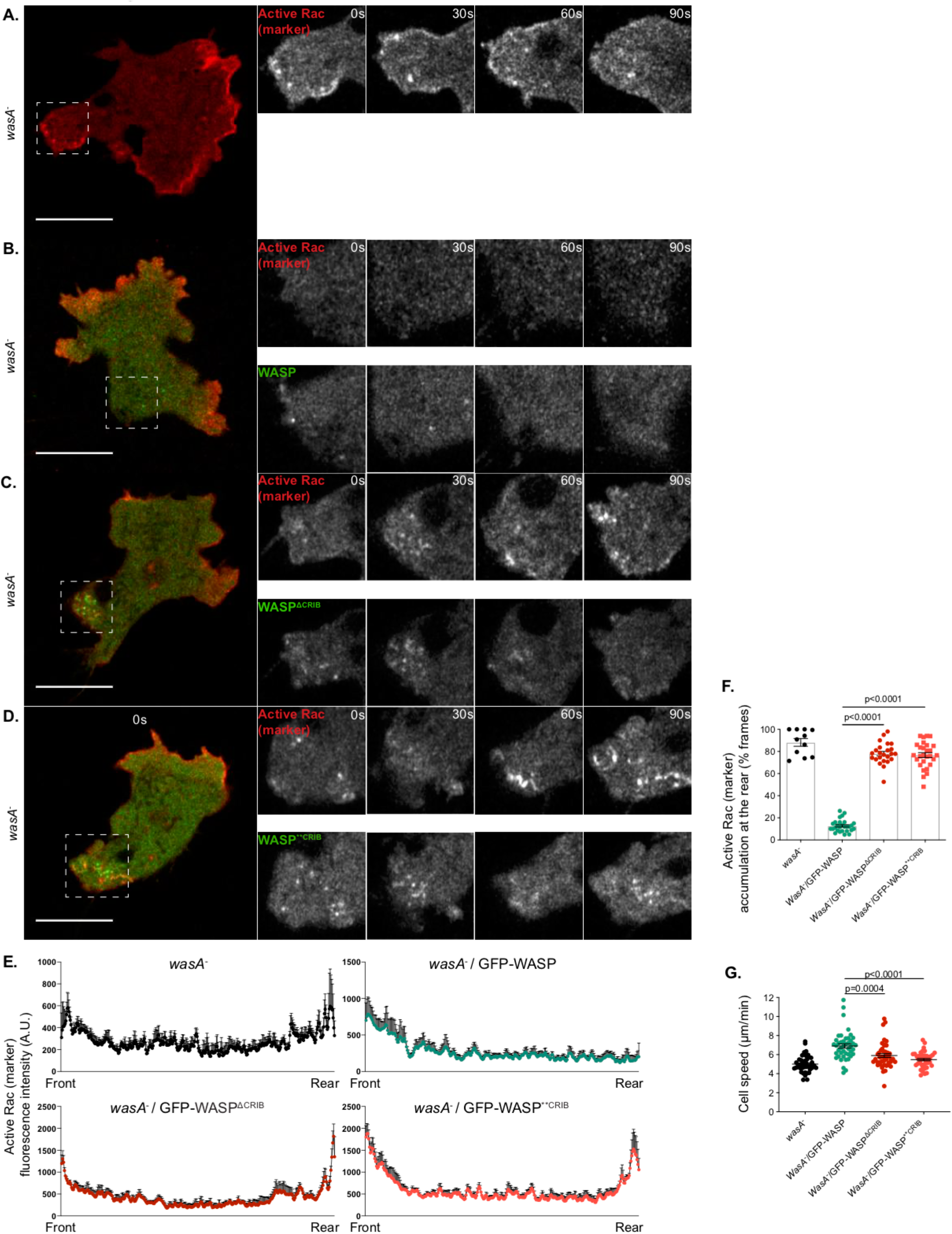
WASP requires a functional CRIB motif to confine active Rac at the leading edge during migration. Airyscan confocal imaging of migrating WASP null cells (*wasA*^-^) expressing an active Rac marker (PakB CRIB-mRFPmars2). Insets show in greater detail the rear of the cell (dashed squares) at different time points. **A.** Active Rac accumulates at the back of *wasA*^-^ cells at all time points. **B.** In cells expressing GFP-WASP no sign of active Rac enrichment at the rear can be detected. In cells expressing GFP-WASP^ΔCRIB^ **(C)** or GFP-WASP^**CRIB^ **(D)**, active Rac can be detected within the enlarged uropod (dashed square). WASP CRIB-mutants generated puncta that cluster at the enlarged rear. Scale bar represents 10 μm. **E.** Fluorescence plots reporting the intensity of the active Rac marker at four time points along a straight line that connects the leading edge to the rear of WASP null cells (*wasA*^-^, top left graph) and WASP null cells expressing either WASP (*wasA*^-^/GFP-WASP top right graph), WASP^ΔCRIB^ (*wasA*^-^/GFP-WASP^ΔCRIB^ bottom left graph), or WASP^**CRIB^ (*wasA*^-^/GFP-WASP^**CRIB^ bottom right graph). **F.** Frequency of active Rac accumulation at the back of WASP null cells and WASP null cells expressing wild type or WASP CRIB mutants. *wasA*^-^ (n= 11 cells): 88.2% ± 3.5; *wasA*^-^/GFP-WASP (n= 28 cells): 12.7% ± 1.0; *wasA*^-^/GFP-WASP^ΔCRIB^ (n= 24 cells): 78.1% ± 2.0; *wasA*^-^/GFP-WASP^**CRIB^ (n= 28 cells): 76.7% ± 2.2; means ± SEM. One-way ANOVA, *wasA*^-^/GFP-WASP vs *wasA*^-^/GFP-WASP^ΔCRIB^: p<0.0001; *wasA*^-^/GFP-WASP vs *wasA*^-^/GFP-WASP^**CRIB^: p <0.0001. **G.** Migratory speed of WASP null cells and WASP null cells expressing wild type WASP or WASP CRIB mutants. *wasA*^-^ (n= 45 cells): 5.0 μm/min ± 0.1; *wasA*^-^/GFP-WASP (n= 44 cells): 6.9 μm/min ± 0.2; *wasA*^-^/GFP-WASP^ΔCRIB^ (n= 44 cells): 5.9 μm/min ± 0.2; *wasA*^-^/GFP-WASP^**CRIB^ (n= 41 cells): 5.5 μm/min ± 0.1; means ± SEM. Kruskal-Wallis test, *wasA*^-^/GFP-WASP vs *wasA*^-^/GFP-WASP^ΔCRIB^: p=0.0004; *wasA*^-^/GFP-WASP vs *wasA*^-^/GFP-WASP^**CRIB^: p<0.0001.

In line with previous reports from *wasA*^-^ cells (Davidson et al., 2018), cells expressing either WASP CRIB mutant also migrate at a significantly reduced speed in comparison to those expressing wild type WASP (Fig. 6G).

Our results show that WASP requires an intact CRIB motif to confine active Rac at the leading edge during migration. CRIB mutant WASPs, even though they localise correctly and activate actin polymerization, cannot restrict Rac nor restore normal cell movement.

### WASP contributes to homeostasis of active Rac levels

We examined whether WASP works locally, to spatially confine active Rac at the front, or has a more global role in Rac regulation.

Taking into account that active Rac is largely membrane-bound (Moissoglu et al., 2006), and that the CRIB-based marker interacts selectively with GTP-bound Rac proteins (de la Roche et al., 2005) but is cytosolic when unbound, then the ratio of RFP-CRIB (active Rac marker) atthe membrane and in the cytosol offers a measure of the total quantity of the active Rac in the cell (Veltman et al., 2012). We therefore measured the relative quantities of RFP-CRIB at the membrane and within the cytosol in cells expressing CRIB-mutated WASPs. As shown in figure 7A and quantified in B, the membrane:cytosol ratio of the active Rac marker is consistently and significantly higher in cells expressing a WASP CRIB mutant in comparison to cells expressing wild type WASP. Thus, Rac activation is generally higher in WASP mutants. We confirmed this result by pulling down the active Rac from cell lysates using GST-CRIB beads (Figure 7C). Although this assay is not quantitative, it clearly shows a higher level of activated Rac in cell expressing WASP CRIB mutants.

**Figure 7.**
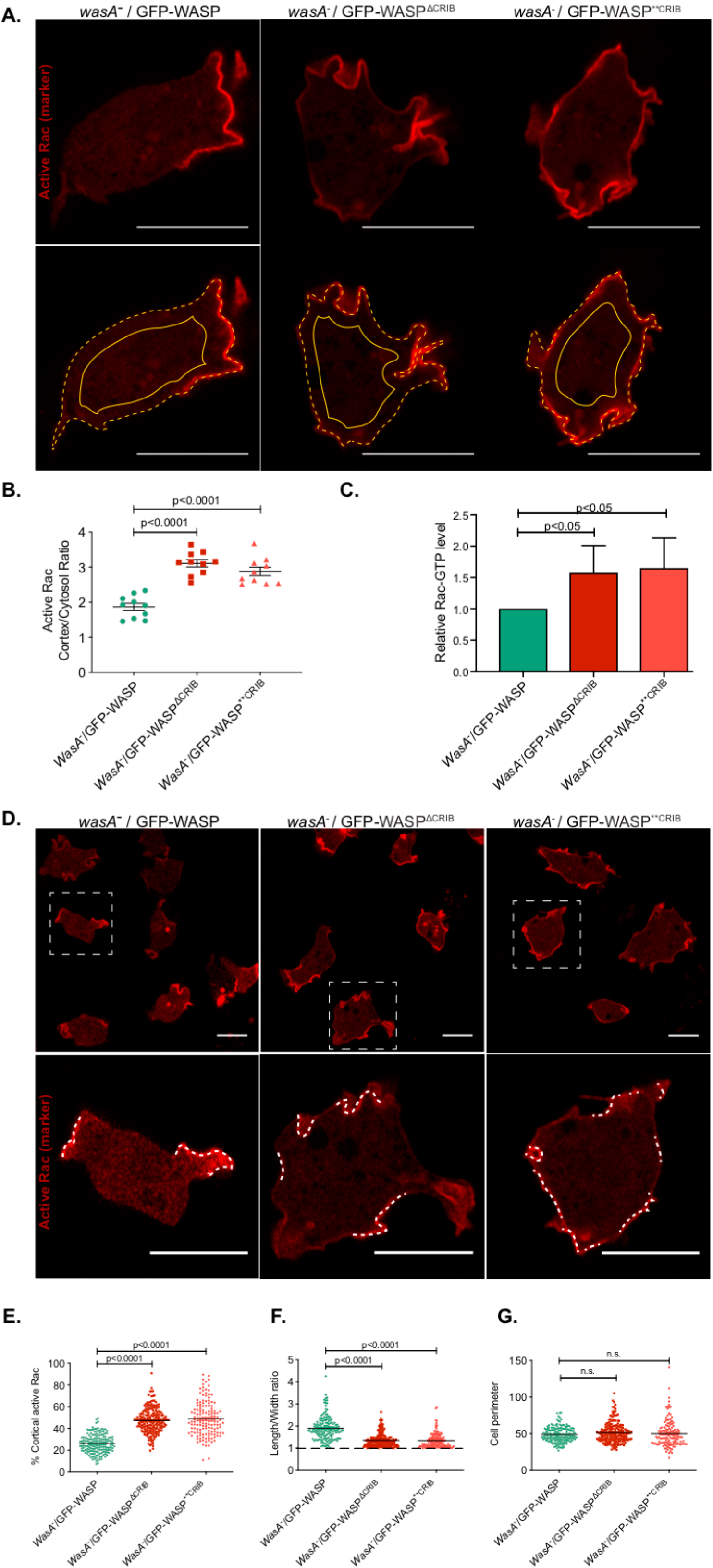
WASP requires a functional CRIB motif to maintain homeostatic levels of membrane-bound active Rac. **A.** Airyscan confocal imaging of WASP null cells (*wasA*^-^) co-expressing an active Rac marker (PakB CRIB-mRFPmars2) and a GFP-tagged WASP (wild type, ΔCRIB, or **CRIB; green channel not shown). Images were acquired consistently through the mid-point of cells. The ratio between the intensity of the active Rac marker along the plasma membrane (dashed line) and the cytosol (solid line) was measured. Scale bars represent 10 μm. **B.** Quantification of the active Rac marker membrane:cytosol intensity ratio from multiple cells as in panel A. *wasA*^-^/GFP-WASP (n= 10 cells): 1.9 ± 0.1; *wasA*^-^/GFP-WASP^ΔCRIB^ (n= 10 cells): 3.1 ± 0.1; *wasA*^-^/GFP-WASP^**CRIB^ (n= 10 cells): 2.9 ± 0.1; means ± SEM. One-way ANOVA, *wasA*^-^/GFP-WASP vs *wasA*^-^/GFP-WASP^ΔCRIB^: p<0.0001; *wasA*^-^/GFP-WASP vs *wasA*^-^/GFP-WASP^**CRIB^: p<0.0001. **C.** Pulldown assay for active Rac levels. Wild type and WASP mutant cells were lysed and active GTP-bound Rac was selectively precipitated using GST-PAK-CRIB immobilised on glutathione agarose. Active Rac was measured by Western blotting using an anti-Rac antibody (Merck Millipore) and fluorescent secondary (n=3 independent experiments; single-tailed t-test). **D.** Airyscan confocal imaging of *wasA*^-^ cells co-expressing an active Rac marker (PakB CRIB-mRFPmars2) and a GFP-tagged WASP (wild type, ΔCRIB, or **CRIB; green channel not shown). Cells expressing wild type WASP (left panel) accumulate active Rac on a small proportion of their plasma membrane (dashed lines within the inset). Cells expressing WASP^ΔCRIB^ or WASP^**CRIB^ (central and right panel respectively) accumulate active Rac in a larger proportion of their plasma membrane (dashed lines within the inset). Scale bars represent 10 μm. **E.** Percentage of plasma membrane enriched in active Rac in cells expressing wild type or CRIB-mutated WASPs. Multiple cells from an experiment as in panel C. *wasA*^-^/GFP-WASP (n= 168 cells): 25.9% ± 0.7; *wasA*^-^/GFP-WASP^ΔCRIB^ (n= 204 cells): 47.3% ± 0.8; *wasA*^-^/GFP-WASP^**CRIB^ (n= 145 cells): 48.8% ± 1.3; means ± SEM. Mann Whitney test, *wasA*^-^/GFP-WASP vs *wasA*^-^/GFP-WASP^ΔCRIB^: p<0.0001; *wasA*^-^/GFP-WASP vs *wasA*^-^/GFP-WASP^**CRIB^ : p<0.0001. **F.** Length to width ratio of cells expressing wild type or CRIB-mutated WASP. Multiple cells from an experiment as in panel C. *wasA*^-^/GFP-WASP (n= 168 cells): 1.9 ± 0.04; *wasA*^-^/GFP-WASP^ΔCRIB^ (n= 204 cells): 1.4 ± 0.02; *wasA*^-^/GFP-WASP^**CRIB^ (n= 145 cells): 1.3 ± 0.03; means ± SEM. Mann Whitney test, *wasA*^-^/GFP-WASP vs *wasA*^-^/GFP-WASP^ΔCRIB^: p<0.0001; *wasA*^-^/GFP-WASP vs *wasA*^-^/GFP-WASP^**CRIB^ : p<0.0001. **G.** Perimeter of cells expressing wild type or CRIB-mutated WASP. Multiple cells from an experiment as in panel C. *wasA*^-^/GFP-WASP (n= 168 cells): 48.8 ± 0.8; *wasA*^-^/GFP-WASP^ΔCRIB^ (n= 204 cells): 51.3 ± 0.9; *wasA*^-^/GFP-WASP^**CRIB^ (n= 145 cells): 49.9 ± 1.5; means ± SEM. Kruskal-Wallis test, *wasA*^-^/GFP-WASP vs *wasA*^-^/GFP-WASP^ΔCRIB^: p=0.56, *wasA*^-^/GFP-WASP vs *wasA*^-^/GFP-WASP^**CRIB^: p = 0.84.

We also examined the extent of active Rac patches. Cells expressing wild type WASP accumulate the active Rac marker on a relatively small portion of their plasma membrane, generally corresponding to macropinocytic cups. By contrast, cells expressing CRIB-mutated WASPs tend to accumulate the active Rac marker in a significantly larger portion of their plasma membrane (Fig. 7D, quantified in E). The increase in plasma membrane enriched in active Rac marker is accompanied by a reduction in cell polarity (Fig. 7F). Indeed, while cells expressing wild type WASP appear relatively elongated even when migrating randomly, those expressing either WASP CRIB mutants have a more rounded morphology, with a length-to-width ratio of nearly one.

No significant difference between the average perimeter of cells expressing wild type or CRIB-mutated WASPs was seen (Fig. 7G), so the more widespread localisation of the active Rac marker in cells expressing WASP CRIB mutants is not due to the availability of more plasma membrane. Together, these results suggest that WASP has a key role in restricting patches of membrane-bound active Rac.

We confirmed the correlation between active Rac levels and the proportion of plasma membrane labelled by the active Rac marker using an inducible dominant-active Rac. Wild type cells expressing the RFP-tagged active Rac marker were transfected with tetracycline-inducible, untagged G12V Rac1A. Induction with tetracycline causes them to accumulate the active Rac marker on a significantly higher proportion of the plasma membrane, often close to 100% (Fig. S3A and B), and become markedly rounder (Fig. S3C). This confirms that WASP relies on a functional CRIB motif to restrict active Rac patches. Altogether, our data show that cells whose WASP lacks a functional CRIB motif, and thus cannot bind Rac, accumulate more active Rac, on a higher portion of the plasma membrane than cells with an intact WASP.

### WASP requires Rac interaction to replace SCAR/WAVE at pseudopods

We have shown that WASP is equally able to trigger actin polymerisation on clathrin pits whether it can or cannot interact with active Rac. Internalisation of CCPs is by far the clearest and evolutionarily conserved role of WASP; however, its behaviour is plastic, and changes in response to different conditions. For instance, WASP can drive extension of protrusions in cells that have lost SCAR/WAVE (Veltman et al., 2012). It remains unclear how cells determine that SCAR/WAVE is inactive, but when it occurs, WASP localises to the leading edge of cells and drives extension of actin-rich protrusions downstream of active Rac1 (Veltman et al., 2012). We therefore explore the possibility that WASP can drive pseudopods extension independently of Rac1, as we have observed during CME.

To address this question, we took advantage of a cell line in which *wasA* was knocked out and the SCAR/WAVE protein was expressed under the control of a tetracycline-inducible promoter (*wasA*^-^ *scrA*^tet^ cells; (Davidson et al., 2018). We co-expressed GFP-WASP (wild type or CRIB mutants) and LifeAct-RFP in *wasA*^-^ *scrA*^tet^ cells, and examined the ability of transfected cells to migrate toward a source of chemoattractant in the presence of tetracycline (when the SCAR/WAVE protein is expressed) or after this was removed (when the only pseudopod-inducer was the exogenously-expressed WASP).

As shown in figure 8A, cells expressing SCAR/WAVE and wild type WASP (*wasA*^-^ *scrA*^tet/ON^/GFP-WASP) migrate efficiently in a gradient of chemoattractant (red line: start point, yellow line: end point); under these circumstances, WASP is detected within F-actin-rich puncta. Once the expression of SCAR/WAVE is turned off (figure 8B), cells expressing wild type WASP (*wasA*^-^ *scrA*^tet/OFF^/GFP-WASP) are still able to move up a gradient of chemoattractant. Under these circumstances, WASP is additionally localised to the leading edge where it drives formation of pseudopods, as previously described (Veltman et al., 2012). Cells expressing WASP^ΔCRIB^ or WASP^**CRIB^ migrate fairly efficiently toward a source of chemoattractant as long as the expression of SCAR/WAVE is maintained (Fig 8C and E, *wasA*^-^ *scrA*^tet/ON^/GFP-WASP^ΔCRIB^ and *wasA*^-^ *scrA*^tet/ON^/GFP-WASP^**CRIB^ respectively). In each case, the mutated WASP is found within puncta that cluster within the enlarged uropod as described earlier. When expression of SCAR/WAVE is turned off, cells expressing either WASP CRIB mutant completely lose the ability to move, in exactly the same way as the cells lacking SCAR and WASP (Davidson et al., 2018), even when driven by a chemoattractant gradient (figures 8D and F, *wasA*^-^ *scrA*^tet/OFF^/GFP-WASP^ΔCRIB^ and *wasA*^-^ *scrA*^tet/OFF^/GFP-WASP^**CRIB^ respectively). Normal, broad pseudopods are no longer generated; they are replaced by obviously inefficient spiky protrusions, which cannot translocate the cell.

**Figure 8.**
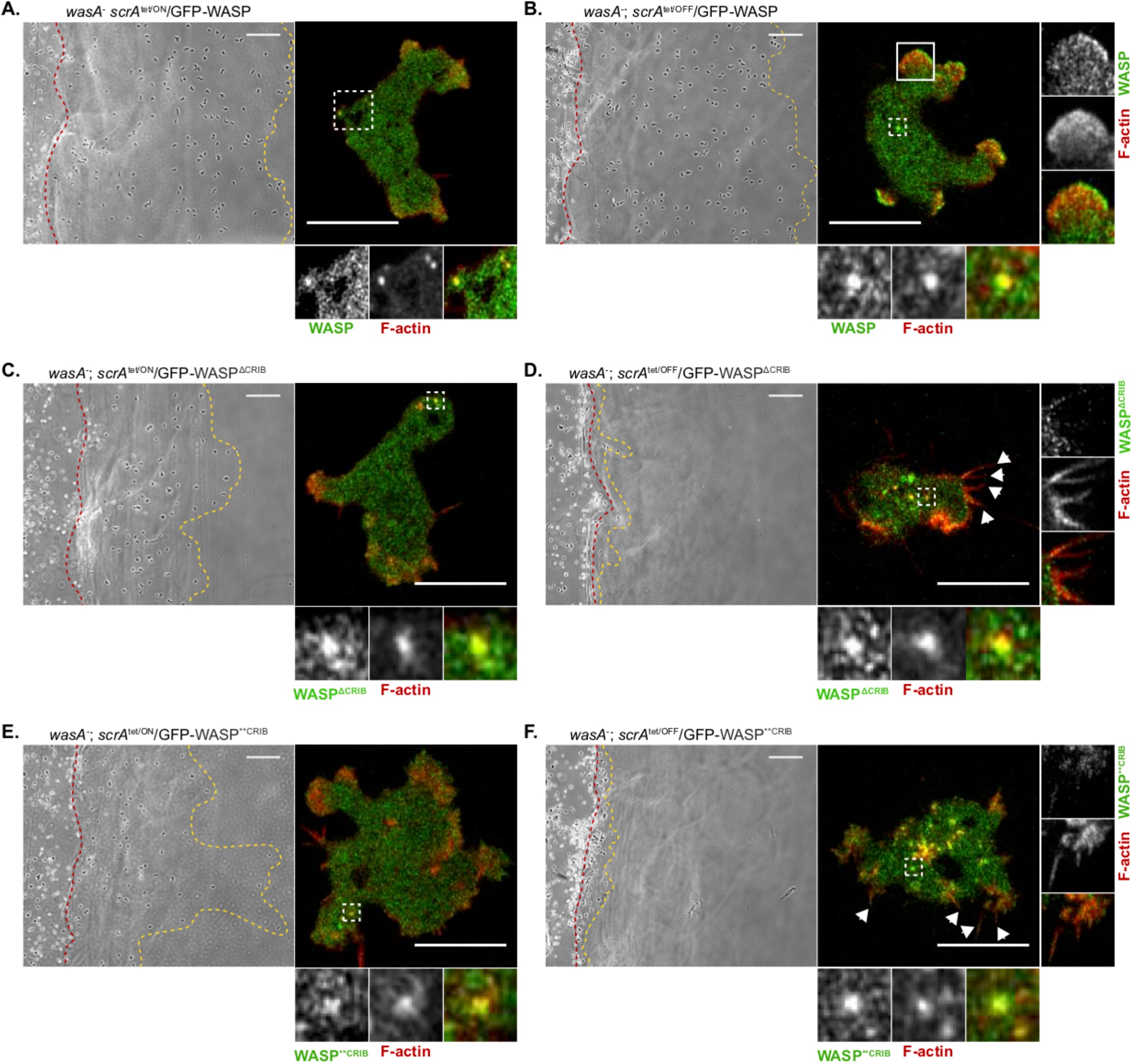
WASP requires a direct interaction with active Rac to drive pseudopod extension in the absence of SCAR/WAVE. **A.** Inducible double null cells expressing GFP-WASP were kept in the presence of tetracycline to maintain SCAR/WAVE expression (*wasA*^-^ *scrA^tet/^*^ON^/GFP-WASP), allowed to migrate for 10 hours toward a chemoattractant and imaged by phase contrast microscopy. Cells migrate efficiently from the starting point (red line) to the end point (yellow line). Scale bar represents 100 μm. As reported on the right-hand side, airyscan confocal imaging shows that GFP-WASP localises to actin-rich puncta at the rear half of the cell (dashed square). Scale bar represents 10 μm. **B.** Inducible double null cells expressing GFP-WASP were kept in the absence of tetracycline for 48 hours to suppress SCAR/WAVE expression (*wasA*^-^; *scrA^tet/^*^OFF^/GFP-WASP), allowed to migrate for 10 hours toward a chemoattractant and imaged by phase contrast microscopy. Cells migrate efficiently from the starting point (red line) to the end point (yellow line). Scale bar represents 100 μm. As reported on the right-hand side, airyscan confocal imaging reveals that GFP-WASP localises to actin-rich puncta at the rear half of the cell (dashed square) as well as at the front (solid square), where it drives pseudopods extension. Scale bar represents 10 μm. **C.** and **E.** Inducible double null cells expressing GFP-WASP^ΔCRIB^ and GFP-WASP^**CRIB^ respectively were grown in the presence of tetracycline to maintain SCAR/WAVE expression (*wasA*^-^; *scrA^tet/^*^ON^/GFP-WASP^ΔCRIB^ and *wasA*^-^; *scrA^tet/^*^ON^/GFP-WASP^**CRIB^ respectively), allowed to migrate for 10 hours toward a chemoattractant and imaged by phase contrast microscopy. Cells are able to migrate from the starting point (red line) to the end point (yellow line), although less efficiently than cells shown in A. Scale bars represent 100 μm. As shown on the right-hand panels, airyscan confocal microscopy shows that GFP-WASP^ΔCRIB^ and GFP-WASP^**CRIB^ generate actin-rich puncta that accumulate at the enlarged rear (dashed squares). Scale bars represent 10 μm. **D.** and **F.** Inducible double null cells expressing GFP-WASP^ΔCRIB^ and GFP-WASP^**CRIB^ respectively were deprived of tetracycline 48 hours to suppress SCAR/WAVE expression, (*wasA*^-^; *scrA^tet/^*^OFF^/GFP-WASP^ΔCRIB^ and *wasA*^-^; *scrA^tet/O^*^FF^/GFP-WASP^**CRIB^ respectively), allowed to migrate toward a chemoattractant, and imaged after 10 hours using phase contrast microscopy. Under these circumstances, cells do not migrate under-agarose, as demonstrated by the substantial proximity of the red and yellow lines (start and end point respectively). Scale bars represent 100 μm. As shown on the right-hand side panels, airyscan confocal imaging reveals that cells expressing a CRIB-mutated WASP are not able to generate pseudopods and form spiky protrusions instead (white arrowheads). The ability of both WASP CRIB-mutant to generate actin-rich puncta is not affected (dashed squares). Scale bars represent 10 μm.

Thus, WASP is able to respond to different stimuli in order to fulfil different cellular roles. Rac activity is not necessary for WASP’s activation in clathrin-mediated endocytosis, but a functional CRIB motif, and thus presumably direct activation by Rac, is absolutely essential for it to stimulate pseudopod formation and leading edge extension in the absence of SCAR/WAVE.

## Discussion

### WASP’s interaction with small GTPases: implications for WASP regulation

Actin polymerization occurs through self-assembly. Once a new filament is initiated, actin monomers contain all the information they need to extend it – but the process must be precisely controlled to function physiologically. One principal way that cells utilise to achieve spatio-temporal control of actin polymerisation is through nucleation promoting factors (NPFs), which determine when and where the Arp2/3 is included into filaments to trigger their branching and growth. Considering how important NPFs are, for processes as diverse as cell movement (Evans et al., 2013; Kunda et al., 2003; Veltman et al., 2012), macropinocytosis (Veltman et al., 2014), vesicular sorting (Carnell et al., 2011; Derivery et al., 2009), mitosis (King et al., 2010), meiosis (Burdyniuk et al., 2018), and DNA repair (Caridi et al., 2018), their control is exceptionally poorly understood. The SCAR/WAVE complex contains five proteins (Eden et al., 2002) and harbours multiple potential phosphorylation and protein-protein interaction sites (Pocha and Cory, 2009), but the relevance of most is largely unknown. Control of the sorting regulators WASH and WHAMM is even less well understood. WASPs, the family that includes mammalian WASP and N-WASP and the singular *Dictyostelium* WASP, have been thought to be an exception. In WASPs, binding of active small GTPases (Cdc42 or Rac), PIP_2_, and/or adaptor proteins has been proposed as essential for activation (Carlier et al., 2000; Miki et al., 1996; Miki et al., 1998; Papayannopoulos et al., 2005; Prehoda et al., 2000; Rohatgi et al., 1999; Tomasevic et al., 2007). However, in this work we show that *Dictyostelium* WASP does not require small GTPase binding or activity for either its localisation or activation.

While this was surprising – a number of reviews describe GTPase binding as an essential step towards both (Kessels et al., 2011; Prehoda and Lim, 2002)– it was not unprecedented. Yeasts and other fungal WASPs have lost their Rac-binding CRIB motifs during evolution, but operate in clathrin-mediated endocytosis by a remarkably similar mechanism (Madania et al., 1999). Furthermore, deletion experiments in *Drosophila* show that WASP can genetically complement a mutant when its CRIB domain is deleted (Tal et al., 2002). Similarly, overexpression of CRIB-deleted N-WASP still causes actin-induced vesicle rocketing in cultured murine fibroblasts (Benesch et al., 2002). The mutation we have developed in this work is a finer tool - it relies on only two conservative substitutions, rather than a deletion, to abrogate the interaction with active Rac – but we show it still allows efficient localisation to clathrin-coated pits and activation of actin polymerisation. Thus, the mechanism by which WASP is activated in living cells is still a surprisingly unsolved conundrum.

### WASP’s role in front–rear polarity

If WASP does not need small GTPases to localise to clathrin-coated pits or to trigger actin polymerisation on puncta, why does it interact with them? CRIB motifs are the best-understood GTPase interaction motifs, and give high specificity for the active (GTP-bound) forms of Rac and Cdc42 (Burbelo et al., 1995). *Dictyostelium* lacks a Cdc42 (Rivero et al., 2001), which first appears just before fungal and metazoan evolutionary lineages divide, but its WASP has a well-conserved CRIB motif which gives strong binding specificity to the active form of Rac1A and C (Han et al., 2006). What are its physiological roles?

The work we describe provides two related answers. Firstly, WASP needs active Rac to replace SCAR/WAVE and drive pseudopods. WASPs do not appear to have a role in pseudopod formation under normal circumstances (Sarmiento et al., 2008), but retain the ability to take over when SCAR/WAVE is absent (Zhu et al., 2016), re-localising to the cell front and driving protrusion (Veltman et al., 2012). It remains unclear how they detect that SCAR/WAVE is lost, but since active Rac is the key specifier of the leading edge, it is unsurprising that WASP requires Rac binding to do this job.

The second role for Rac binding to WASP is more subtle. When there is no WASP, cells lose polarity in a curious and idiosyncratic way – as they migrate, they generate pseudopods as usual at the front, but also carry a substantial and structurally complex assembly at the rear that slows and depolarizes them. This aggregate contains active Rac and continuously recruits new SCAR/WAVE and thus Arp2/3 complex and actin. This is surprising for at least two reasons. First, it highlights our poor understanding of the cell’s uropod, which may simplistically and wrongly be seen as a black hole where unnecessary migratory components are passively drawn. Second, it underscores our incomplete knowledge of WASP’s roles and subcellular dynamics.

Here we propose a novel role for WASP as a homeostatic regulator of active Rac. We envision a model whereby in addition to triggering actin polymerisation at CCPs, WASP is also capable of removing active Rac from the plasma membrane. This will obviously occur faster when GTPases are excessively activated, and slower when there is little GTPase activation, so it will promote homeostasis of small GTPase activation. Similarly, CME occurs at a greater rate at the rear of migrating cells, so active Rac will be removed faster there. This will help maintain the front enrichment of Rac in normal cells, and explain the disproportionate accumulation at the rear in WASP mutants.

### WASP and CRIB motifs: “the cowl does not make the monk”

When WASP was first discovered, the presence of a CRIB motif led researchers to conclude that its activity was dependent on small GTPases (Miki et al., 1996). This now seems to be an oversimplification. Cells utilise WASP for multiple processes, and we have shown here that the importance of small GTPases for WASP to fulfil its different roles is context-dependent. On one hand, WASP does not rely on the availability of (nor on the direct interaction with) active Rac to recruit Arp2/3 complex, drive actin polymerization, or promote clathrin-mediated endocytosis. On the other hand, WASP depends on a direct, CRIB-mediated, interaction with active Rac to drive pseudopod extension in the absence of SCAR/WAVE. Therefore, WASP’s requirement for active Rac appears to be a context-dependent feature, not an absolute requirement as initially thought. In the broader context, this means that inhibiting Rac may not of itself compromise WASP functionality. WASP has many physiological roles (for example in invadopodia formation), and its CRIB motif is likely to be important for a subset of other processes that we have not explored.

## Materials and Methods

### Cell culture and transfection

*Dictyostelium* wild type (*wasA*^+^), *wasA* knockout (*wasA*^-^), and inducible double null cells were grown on Petri dishes in HL5 supplemented with vitamins and micro-elements (Formedium™). Prior to transfection, cells were resuspended in E-buffer (10 mM KNaPO_4_, pH 6.1, 50 mM sucrose), incubated with DNA of interest, and elecroporated at 500 V using the ECM 399 system (Harvard Apparatus U.K.). Transfected cells were then transferred in Petri dishes containing medium, and selected 24 hour later by addition of 50 μg/ml hygromycin.

### Protein purification

E. coli BL21 cells were transformed with a pGEXRac1A-encoding vector by heat-shock at 42 ° C (plasmid was a gift from A. Kortholt, University of Groningen, the Netherlands), and plated overnight on ampicillin-containing SM plates. One colony was transferred to a tube containing LB medium (1% Bacto-tryptone, 0.5% Bacto-yeast extract, 17 mM NaCl, pH 7) plus ampicillin, and kept in shaking condition (200 rpm) at 37° C until it reached the OD_600_ of 2. The bacterial suspension was then transferred in a larger volume of LB medium plus ampicillin, and kept overnight in shaking conditions (100 rpm) at room temperature. Once the bacterial suspension reached the OD_600_ of 0.4, 500 μM IPTG (Isopropyl β-D-1 thiogalactopyranoside) was added to yield the expression of GST-Rac1A, and bacteria were kept overnight in shaking conditions (100 rpm) at room temperature. Bacteria were then spun down using the Avanti J6-MI centrifuge (Beckman Coulter) at 4° C 3000 rpm for 20 minutes, and lysed using a detergent-based lysis buffer (1% Triton X-100, Halt™ Protease Inhibitor Cocktail, 1mM DTT).

The lysate was sonicated 10 times at the maximum power with 10 seconds interval between cycles using a Soniprep 150 (MSE), and spun down at 4° C for 10 minutes at 12096×g using an Avanti J-25 centrifuge (Beckman Coulter). The sonicated lysate was incubated for 2 hours at 4 ° C with beads (Glutathione High Capacity Magnetic Agarose Beads, Sigma-Aldrich) previously washed in Rac buffer (50 mM Tris HCl pH 7.5, 100 mM NaCl, Halt™ Protease Inhibitor Cocktail, 1mM DTT). The beads were then washed in Rac buffer and incubated with 1 mM GTPγs (Guanosine 5ʹ-[β,γ-imido]triphosphate trisodium salt hydrate, Sigma-Aldrich) overnight at 4 ° C in the presence of 15 mM EDTA and 100 units of CIP. MgCl_2_ was ultimately added to the tube at the final concentration of 60 mM in order to close the Rac’s nucleotide-binding pocket.

### GST-Rac pull-down assay and immunoblotting

*Dictyostelium* cells were resuspended in lysis buffer (50 mM Tris HCl pH 8, 100 mM NaCl, 30 mM MgCl_2_, 0.1% Triton X-100, Halt™ Protease Inhibitor Cocktail, 1mM DTT) and the resulting lysate added to the tube containing GTPγs-Rac1A-loaded beads for 1 hour at 4° C. The tubes were placed on a magnet and washed twice in washing buffer (50 mM Tris HCl pH 8, 100 mM NaCl, 30 mM MgCl_2_, Halt™ Protease Inhibitor Cocktail, 1mM DTT). Beads were then washed twice with 1 ml of washing buffer, resuspended in NuPAGE™ LDS Sample Buffer (ThermoFisher) and incubated at 100 ° C for 5 minutes. Proteins contained in the eluted fraction were separated using NuPAGE™ 4-12% Bis-Tris protein gels (ThermoFisher). Protein were transferred on nitrocellulose membrane (Amersham^™^ Protran^®^, Merck) at 80 V for 1 hour. The membrane was blocked for 1-hour in the presence of 5% semi-skimmed milk diluted in TBS. The blocked membrane was incubated overnight at 4° C with the desired primary antibody. The membrane was washed three times with TBST and incubated with the appropriate secondary antibody diluted in 5% BSA in TBST. The membrane was washed three times with TBST, scanned using Odyssey CLx (LI-COR) and analysed using Image Studio software (LI-COR).

### Active Rac pull-down assay

Active Rac levels were determined by pull-down assay using GST-PAK-CRIB immobilised to Glutathione beads according to manufacturer’s instructions (Cytoskeleton, Inc, Active Rac Kit). Briefly, *wasA-* cells expressing GFP-WASP (*wasA*-/GFP-WASP) or respective CRIB mutants (*wasA-*/GFP-WASP^**CRIB^ and *wasA-*/GFP-WASP^ΔCRIB^) were collected from a confluent 30 cm petri dish and resuspended in lysis buffer supplemented with HALT^TM^ EDTA-free Protease Inhibitor Cocktail at 4 ° C. After a brief clarification at 10,000 g for a minute, the supernatant is loaded into GST-PAK-CRIB beads in eppendorf tubes and incubated for 1 hour at 4 ° C with gentle rotation. The washed beads were then resuspended in 2x sample buffer and boiled for 3 minutes at 95 ° C. Proteins contained in the eluted fractions were separated using NuPAGE™ 12% Bis-Tris protein gels (ThermoFisher) and transferred into nitrocellulose membrane as described previously. The membrane is then blocked for 30 minutes in the presence of 5% semi-skimmed milk diluted in TBS. The blocked membrane was incubated overnight at 4 ° C with a mouse monoclonal anti-Rac antibody (1/500 dilution). The membrane was washed once with TBST and incubated with appropriate secondary antibody diluted in 5% BSA in TBST. After three washes, the membrane was scanned using Odyssey CLx (LI-COR) and analysed using Image Studio software (LI-COR). Wild type cells expressing G12V Rac1A under a tetracycline-inducible promoter was used a positive control in the above experiments. In short, cells were treated with tetracycline to the final concentration of 10 μg/ml to induce Rac G12V expression. Cells were harvested 2-3 hours post-induction and processed as described above. The levels of active Rac is quantified using Image Studio Lite^TM^ and relative active-Rac levels are then calculated and plotted using Prism 7.

### Microscopy

Super-resolution microscopy was adopted to investigate the ability of WASP CRIB mutants to localise to clathrin-coated pits and to recruit the Arp2/3 complex. Images were acquires using a Zeiss LSM880 equipped with a 63x/1.40 NA objective. GFP was excited at 488 nm, RFP at 561 nm. Images were acquired using the ZEN imaging software every 1 or 2 seconds.

TIRF microscopy was utilised to monitor the dynamics of clathrin, WASPs, Arp2/3 complex and actin on the ventral surface of the cells. Images were acquired using a modified Nikon Eclipse TE 2000-U microscope equipped with a photometrics Evolve 512 camera and a DualView DV2 emission splitter. GFP and RFP were excited using 473 nm and 561 nm wavelengths respectively. A 100x/1.40 NA TIRF objective was used. Images were acquired every 1 or 2 seconds using the MetaMorph software.

Phase contrast microscopy was performed to test the ability of cells expressing WASP CRIB-mutants to migrate following loss of SCAR/WAVE. Images were acquired using a Nikon ECLIPSE TE2000-E inverted microscope with a 10x/0.30 NA objective and equipped with a QImaging Retiga EXi digital camera.

Spinning disk confocal microscopy was utilised to monitor WASP dynamics upon expression of dominant active Rac. Images were acquired using a Nikon Ti-E inverted microscope equipped with a Yokogawa CSU-X spinning disc confocal unit in combination with a High resolution Andor Neo sCMOS camera. A 100x/1.4 NA objective was used. GFP was excited at 488 nm, RFP at 561 nm. Images were acquired using the Andor IQ 2 software.

### Chemotaxis assay

The under-agarose chemotaxis assay (Laevsky and Knecht, 2001) was used to investigate the ability of cells expressing no WASP or WASP CRIB mutants to confine active Rac at the leading edge during migration, as well as the ability of inducible double null cells expressing WASP CRIB mutants to generate pseudopods following loss of SCAR/WAVE. A 50mm glass bottom dish (MatTek) was first treated with 10 mg/ml BSA to reduce the resistance encountered by cells crawling under agarose. Seakem GTG agarose was dissolved to the final concentration of 0.4% in LoFlo (Formedium™), a low fluorescence medium that improves the quality of imaging. Dissolved agarose was cast on a pre-treated glass bottom dish and allowed to set. Once set, the agarose was cut using a scalpel so to create two wells separated by a 5 mm bridge; this was gently wriggled loose to facilitate cells’ crawling. Cells of interest were resuspended in LoFlo, counted using CASY® Model TT Cell Counter (Innovatis), and diluted to a final concentration of 5×10^5^ cells/ml. 200 μl of cell suspension was seeded on the left well created within the agarose layer. The other well was filled with 100 μM folic acid diluted in LoFlo. A square coverslip was carefully lowered down in order to cover both wells and prevent evaporation; the glass bottom dish was thereafter incubated at 22° C. Imaging was performed 3-10 hours after setting up the assay.

### Rac inhibitor treatment

Cells expressing a red-tagged active Rac marker (PakB CRIB-mRFPmars2) or a GFP-tagged WASP were resuspended in LoFlo, and 10^5^ cells seeded on a borosilicate glass 8 well chamber (ThermoFisher). Cells were allowed to adhere and then treated with various concentrations of the Rac inhibitor EHT1864. Images were acquired prior to addition of the inhibitor (to monitor the dynamics of the probe in the presence of active Rac), during addition of the drug (to visualise its most acute effects) and for 10-12 minutes after treatment (to verify whether cells could recover).

### Induction of G12V Rac1 expression

Cells expressing G12V Rac1A under a tetracycline-inducible promoter along with GFP-WASP or RFP-tagged active Rac marker were resuspended in LoFlo and seeded to a final concentration of 5×10^5^ cells/ml on a borosilicate glass 8 well chamber (ThermoFisher) or on a 35 mm glass bottom dish (MatTek). Once adhering, cells were treated with tetracycline to the final concentration of 10 μg/ml to yield G12V Rac1A expression. Cells were images from the moment of tetracycline addition up to 2-3 hours post-induction.

### Image processing

Images acquired using phase contrast, TIRF or confocal microscopes were exported as .TIFF files, while those acquired using the super-resolution confocal were exported as .CZI files. Images were all imported into Fiji software (Schindelin et al., 2012) and processed as required, including cropping and linear brightness/contrast adjustment. TIRF images were processed using the Windowed-Sinc Filter (Kunito Yoshida, Department of Biological Sciences, Imperial College London). Clathrin, WASP and Arp2/3 complex *puncta* dynamics was monitored using the MTrackJ plugin (Meijering et al., 2012). The same plugin was utilised along with a Chemotaxis Tool plugin (Gerhard Trapp and Elias Horn, ibidi GmbH) to measure speed and directionality of migrating cells. Images acquired using the super-resolution confocal microscope were subjected to Airyscan processing and to a 2D automatic deconvolution.

### Statistical analysis

Unpaired t-test, Mann-Whitney test, one-way ANOVA with a Kruskal-Wallis multiple comparison test or a Tukey’s multiple comparisons test were performed to generate p values and test for statistical significance.

## Online supplemental material

**Figure S1.**
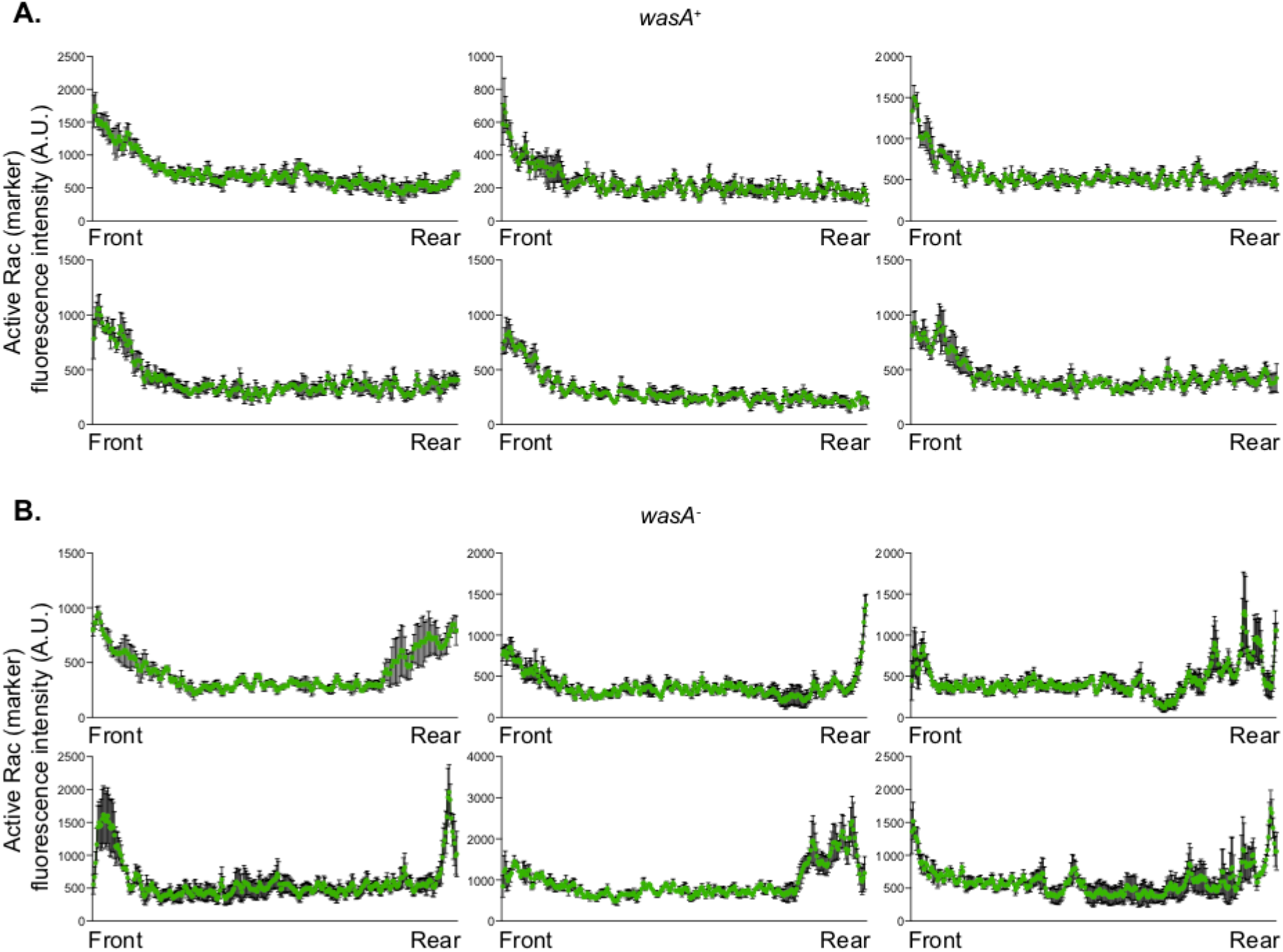
shows additional fluorescence plots reporting the intensity of the active Rac marker along a straight line connecting the front to the rear of wild type (panel A, *wasA*^+^) and WASP null (panel B, *wasA*^-^) cells.

**Figure S2.**
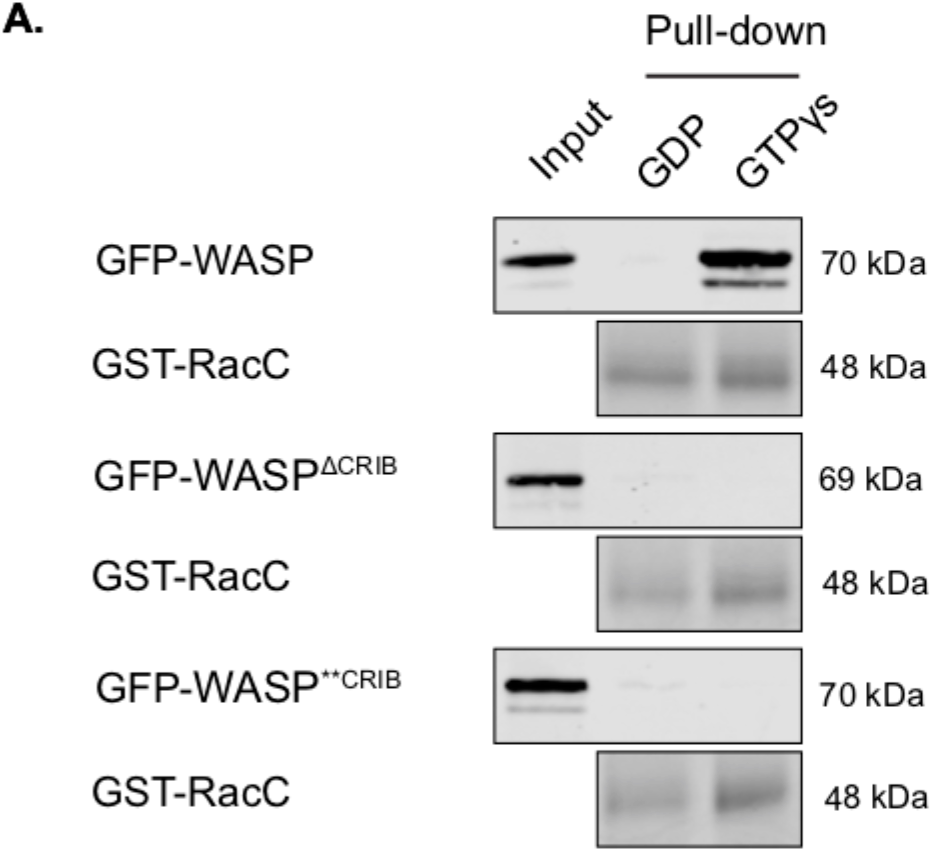
shows the compromised ability of CRIB-mutated WASPs to interact with active RacC, as tested by pull-down assay.

**Figure S3.**
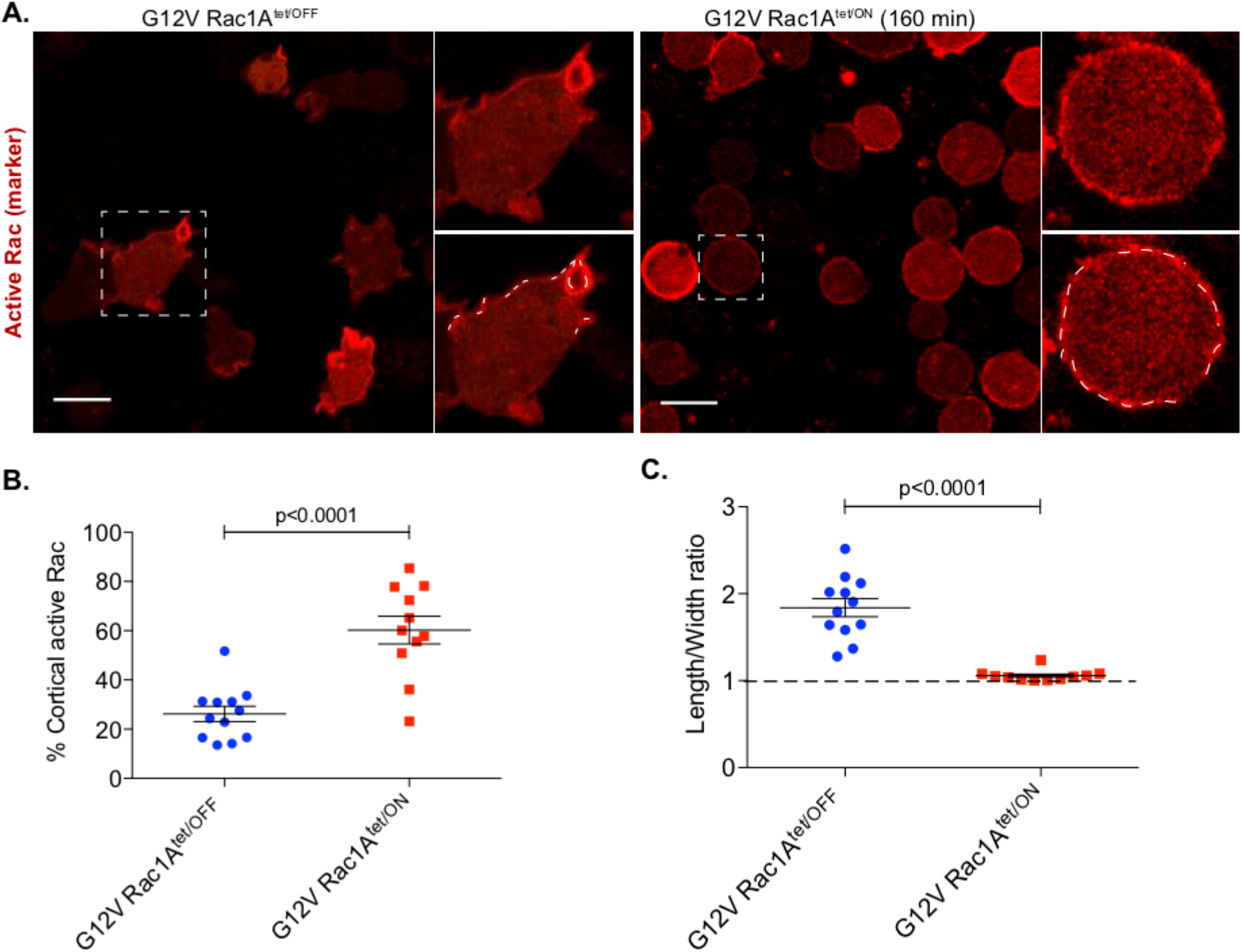
shows the correlation between an increase in the cellular levels of dominant active (G12V) Rac1 and the percentage of plasma membrane labelled by the active Rac marker (panel B) and the ratio between length and width of the cells (panel C).

## Acknowledgements

We thank Margaret O’Prey and Beatson Advanced Imaging Resource (BAIR) staff for assistance with microscopy, Stephen Barratt for assistance with cloning of the CRIB mutants, and Douwe Veltman for providing some plasmids. Funding was provided by Cancer Research UK. Sequences and gene expression data were provided by Dictybase. We are very grateful to members of the Insall and Machesky Labs for helpful comments on the manuscript. The authors declare no competing financial interests.

